# The mitochondrial *orf125* affects male fertility of *Solanum tuberosum* (+) *S. commersonii* somatic hybrids and participates in the onset of “Tuberosum”-Type CMS and evolution of common potato

**DOI:** 10.1101/2024.09.25.614866

**Authors:** Rachele Tamburino, Nunzio D’Agostino, Gaetano Aufiero, Alessandro Nicolia, Angelo Facchiano, Deborah Giordano, Lorenza Sannino, Rosa Paparo, Shin-Ichi Arimura, Nunzia Scotti, Teodoro Cardi

## Abstract

Cytoplasmic male sterility (CMS) is an agronomically significant trait and a powerful tool to study interactions between nuclear and cytoplasmic genomes. In this study, the chondriomes of two isonuclear male-fertile and sterile somatic hybrids (SH9A and SH9B, respectively) between the common potato (*Solanum tuberosum* Group *Tuberosum*, *tbr*) and the wild species *S. commersonii* (*cmm*), were sequenced and compared to those of parental species to identify mitochondrial genes involved in the expression of male sterility. A putative novel gene (*orf125*) was found only in *tbr* and in male-sterile hybrids. Two approaches, a physical or functional deletion of *orf125* by mtDNA editing in SH9B and its allotopic expression in SH9A, clearly demonstrate that *orf125* affects male fertility. To trace the origin of *orf125* and hypothesize its role in the evolution of common potato, we searched it in *tbr* varieties, tuber-bearing potato relatives and other Solanaceae. The organization of the mitochondrial genome region implicated in CMS remained consistent across all common potato accessions in GenBank. An identical *tbr* copy of *orf125* was also detected in all six accessions belonging to the *S. berthaultii* complex (*ber*) analyzed. Such findings corroborate the hypothesis that *ber* accessions with T/β cytoplasm crossed as female with Andean potato (*S. tuberosum* Group *Andigenum*, *adg*), giving rise to the differentiation of the Chilean potato (*S. tuberosum* Group *Chilotanum*), and highlights the origin of mitochondrial factors contributing to genic-cytoplasmic male sterility in *tbr* x *adg* (or some wild species) hybrids.

## Introduction

Cytoplasmic male sterility (CMS) has been identified in over 150 species and can co-exist with hermaphroditism in certain natural populations (gynodioecious species). It serves as a valuable tool to exploit hybridization and heterosis in numerous crops, and study interactions between nuclear and cytoplasmic (mitochondrial) genomes (Budar et al., 2003; Chen and Liu, 2014; Kim and Zhang, 2018; Xu et al., 2022; Kitazaki et al., 2023). CMS can arise spontaneously or following intra-, or more commonly, inter-specific hybridization. This typically occurs due to the expression of specific open reading frames (*S-orfs*), which result from rearrangements in mitochondrial DNA that give rise to entirely novel sequences or to chimeric constructs containing fragments of mitochondrial genes, primarily those encoding subunits of electron transport chain complexes, along with other sequences. Although the sequences of *S-orfs* generally lack conservation, they often feature a transmembrane domain (Hanson and Bentolila, 2004; Chen and Liu, 2014; Kim and Zhang, 2018; Xu et al., 2022; Kitazaki et al., 2023).

Inter-specific somatic hybrids between the common potato (*Solanum tuberosum* Group *Tuberosum, tbr*) and the wild incongruous species *S. commersonii* (*cmm*) were largely male-sterile (Cardi et al., 1993; Cardi, 2001). They showed early degeneration of the tapetum and arrest of meiosis at the conclusion of the reductional phase (Conicella et al., 1997), like other CMS systems and cross combinations involving *tbr* and wild species such as *S. acaule* or *S. curtilobum* (Lamm, 1945; Lamm, 1953; Chen and Liu, 2014; Kitazaki et al., 2023). Remarkably, within one population, an exceptional male-fertile hybrid emerged (Cardi et al., 1993). Male fertility/sterility phenotypes exhibited maternal inheritance, with sterility being partially restored through crossing male-sterile hybrids with a genotype carrying known restorer genes (Iwanaga et al., 1991; Bastia et al., 1999; Cardi et al., 1999). In addition, while male-sterile somatic hybrids predominantly showcased restriction fragment length polymorphisms (RFLPs) derived from the cultivated parent (*tbr*), the male-fertile hybrid displayed a mitochondrial genome more closely resembling, although not entirely identical to, that of *cmm* (Cardi et al., 1999). Finally, parallel work with hybrids resulting from reciprocal sexual crosses between diploid *tbr* and tetraploid (or *2n* pollen producing) *cmm* showed that hybrids were male-sterile when *tbr* was used as the female parent, but male-fertile in the opposite cross direction (Novy and Hanneman, 1991; Carputo et al., 1995). Compounding such information, we hypothesized that the male sterility observed in *tbr* (+) *cmm* somatic hybrids could arise from the interplay between nuclear and mitochondrial genes inherited from *cmm* and *tbr*, respectively. We also speculated that the restoration of male fertility in the single male-fertile hybrid SH9A could depend on the loss of putative mitochondrial genes involved in these interactions. This loss may result from chondriome rearrangements or recombinations following protoplast fusion (Cardi et al., 1999).

Numerous studies primarily focusing on nuclear and plastid DNA suggested that modern potato varieties (*tbr*), bred in North America, Europe, and other parts of the world, mostly derive from the long-day adapted Chilean potato (*S. tuberosum* Group *Chilotanum*) (Spooner et al., 2007). The latter likely resulted from an interspecific cross between a species of the *S. berthaultii* complex, which provided the cytoplasmic genomes, and a tetraploid accession of the short-day adapted Andean potato (*adg*) (Spooner et al., 2007). Within cultivated Andean potatoes (*S. tuberosum* Group *Andigenum*), tetraploid accessions displayed five primary chloroplast types (A, S, C, W, T), exhibiting a gradual variation from the Northern to the Southern Andean regions. Conversely, *S. tuberosum* Group *Chilotanum* predominantly featured the T-type (Hosaka and Hanneman, 1988). On the other hand, limited information is available on mitochondrial DNA diversity in *Solanum spp.* and its potential role in the evolution of *S. tuberosum*, although an attempt has been made to categorize also potato chondriomes into five different types (α, β γ, 8, ι:) (Lössl et al., 1999).

Three cytoplasm types – denominated on the basis of plastome and chondriome composition T/β (commonly found in *tbr*), W/γ (introgressed from *S. stoloniferum*) and W/α (D-type, derived from *S. demissum*) - participate in interactions that often lead to male sterility (Anisimova and Gavrilenko, 2017). The T/β cytoplasm, also recognized as “T”, “Chilean” or “Tuberosum”-type, predominates in most common potato varieties (Hosaka and Sanetomo, 2012; Sanetomo and Gebhardt, 2015) and triggers male sterility when the common potato is crossed as the female parent with *adg* or some wild species as male counterparts (Hermundstad and Peloquin, 1985; Grun, 1990; Anisimova and Gavrilenko, 2017; Goryunova et al., 2023). However, despite the well-established understanding of the cytoplasmic composition of *Solanum* species and common potato genotypes, as well as the compatibility between nuclear and cytoplasmic genomes in interspecific crosses, molecular information regarding the genes responsible for male sterility expression remains limited. Only recently, has a mitochondrial region potentially associated with W/γ-type CMS been proposed, but a sterility inducing *orf* has not yet been identified (Sanetomo et al., 2022).

In tuber-bearing *Solanum* species, nuclear-cytoplasmic male sterility can play a role in maintaining species integrity within sympatric *Solanum* spp. (Camadro et al., 2004). Furthermore, it is a cornerstone for developing innovative breeding strategies centered around True Potato Seed (TPS) reproduction and the development of heterotic F_1_ hybrids (Jansky et al., 2016; Bradshaw, 2022). Finally, the abundance of numerous related cross-compatible species that are amenable to biotechnological approaches highlights their potential as model systems for studying nuclear-cytoplasmic interactions and CMS. Therefore, deciphering the genes that control nuclear-cytoplasmic interactions leading to CMS in *Solanum* spp. holds significant implications not only for biological and evolutionary investigations but also for enhancing genetic diversity and developing innovative breeding strategies in potato cultivation. In this study, complete mitogenome sequencing and multiple analytical approaches, including mtDNA editing and allotopic expression, were employed to identify a candidate mitochondrial gene present exclusively in *tbr* and male-sterile somatic hybrids, and link it to the expression of CMS. Additionally, the validated *S-orf* was searched in *tbr* varieties, tuber-bearing potato relatives and other Solanaceae to trace its origin and hypothesize its broader role in *tbr* evolution and in the expression of “Tuberosum”-type nuclear-cytoplasmic male sterility across potato varieties.

## Results

### Mitochondrial genome rearrangements in male-sterile and fertile somatic hybrids reveal that the tbr-derived orf125 is present only in male-sterile hybrids

The mitochondrial genomes of two isonuclear tetraploid (2*n* = 4*x =* 48) *somatic* hybrids, differing at the phenotypic level only for showing male fertility (SH9A) or male sterility (SH9B), have been sequenced and compared with those of parental species (*cmm* and *tbr*). The mitochondrial genome of the male-sterile hybrid SH9B is fragmented into three molecules 313767, 111810 and 48452 bp long, respectively. It includes 101 non-redundant genes classified as follows: 37 protein coding genes, 2 pseudogenes, 41 open reading frames (ORFs), 3 ribosomal RNAs (rRNAs) and 18 transfer RNAs (tRNAs). Similarly, the mitochondrial genome of the male-fertile hybrid SH9A is fragmented into four molecules with lengths of 251363, 109928, 49622 and 48445 bp. The total number of non-redundant genes is 94, including 37 protein coding genes, 1 pseudogene, 35 ORFs, 3 rRNAs, and 18 tRNAs (Table S1).

The mitogenomes of the two hybrids were then compared with those of the two parents, revealing 23 syntenic blocks ranging from ∼6 kb to 90 kb. Syntenic block 1 was repeated three times in *S. commersonii* and twice in *S. tuberosum* and hybrids. Syntenic block 2 appeared twice in *S. tuberosum* and hybrids. Blocks 3 to 9 were repeated twice in *S. commersonii*. Blocks 18 to 21 were either absent or incomplete (not aligned in their entire length) in *S. commersonii.* Finally, blocks 22 and 23 were missing in *S. commersonii* and SH9A (Fig. 1, Table S2). The comparative analysis clearly highlighted that the SH9B mitogenome closely resembles that of *tbr*, while SH9A exhibits multiple rearrangements compared to both parental species.

**Figure 1.**
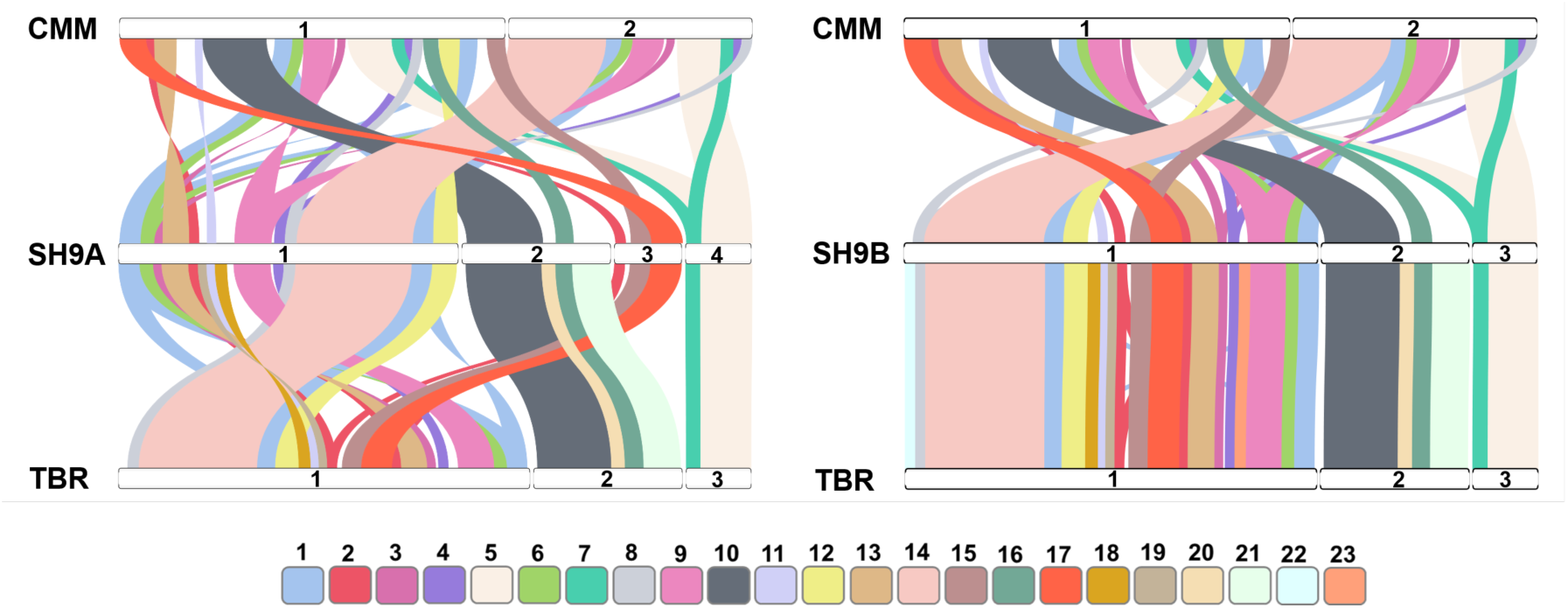
Patterns of synteny between the mitochondrial sequences of the parental species and of SH9A and SH9B somatic hybrids (left and right, respectively). Chromosomes (horizontal bars labeled with numbers) of hybrids, *Solanum commersonii* and *S. tuberosum* cv. Désirée are in the middle, at the top and at the bottom, respectively. SH9A: 1, ON682437; 2, ON682438; 3, ON682439; 4, ON682440. SH9B: 1, ON009139; 2, ON009140; 3, ON009141. *S. commersonii*: 1, MF989960; 2, MF989961. *S. tuberosum* cv. Désirée: 1, MN104801; 2, MN104802; 3, MN104803. Colored ribbons connect the synthetic blocks to each other, each identified by a number and a specific color. A twist in the ribbon indicates an inversion.

Previously, the presence and expression of several *orfs* were assessed in SH9B, SH9A, and the two parental species (Tamburino et al., 2019). PCR and RT-PCR analyses revealed that *orf125a* in syntenic block 14 (hereafter referred to as *orf125*) was present and expressed in flower buds of all five male-sterile hybrids and the cultivated parent *tbr*. Conversely, *orf125* was absent in the male-fertile hybrid SH9A and the wild parent *cmm* (Fig. 2A, B). Further investigations using a custom antibody against ORF125, detected the protein exclusively in the male-sterile hybrid, particularly in anthers from flower buds less than 3 mm (Fig. 2C).

**Figure 2.**
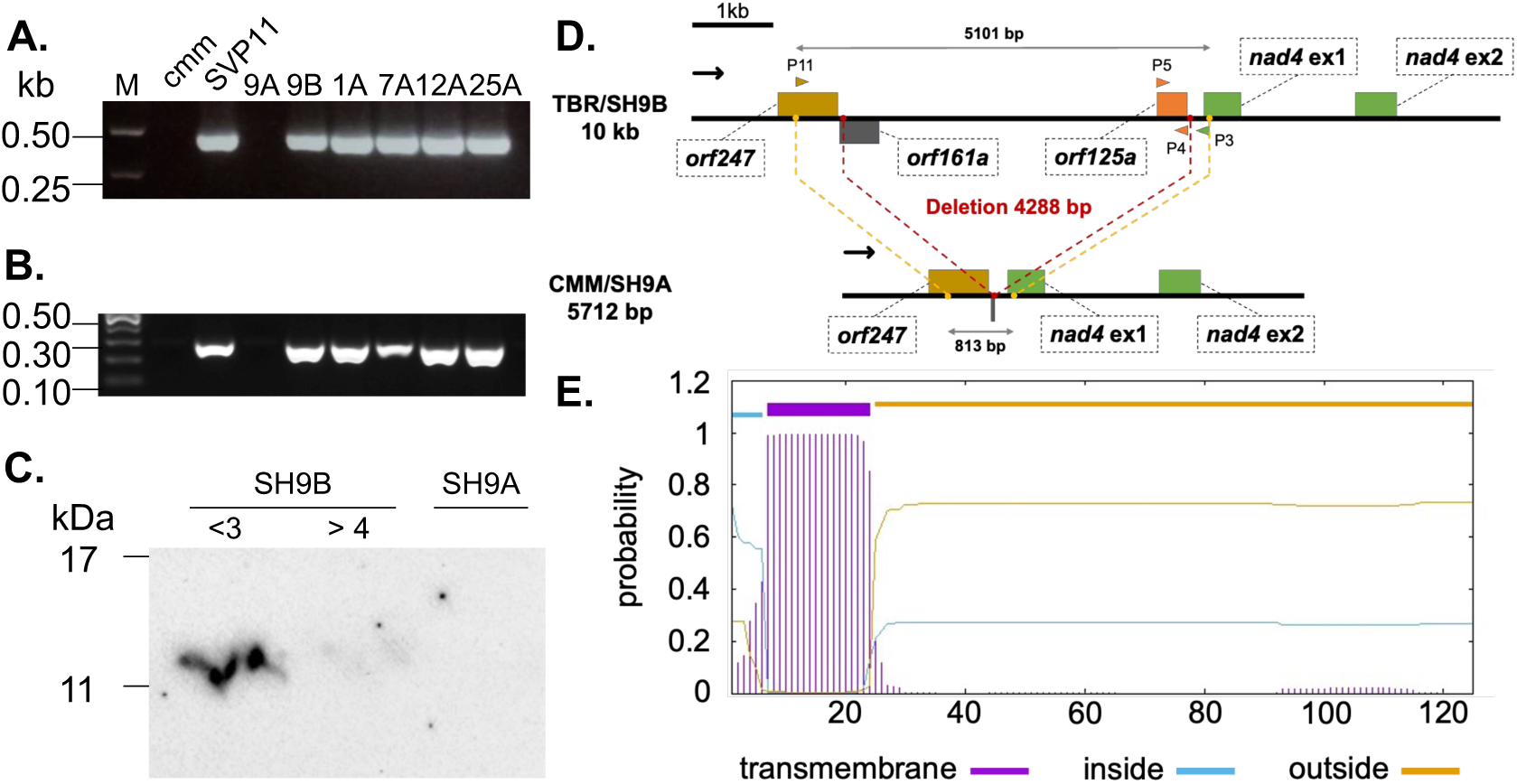
Characterization of *orf125* identified through *in silico* analysis of mtDNA sequences from SH9A, SH9B, *S. commersonii* and *S. tuberosum* cv. Désirée. **A.** PCR results in *S. commersonii* (*cmm*), *S. tuberosum* (SVP11), and a sample of somatic hybrids (9A, male-fertile; 9B and 1A-25A, male sterile) with P4-P5 primers (see Table S5). **B.** RT-PCR results of RNA isolated from flower buds of *cmm*, SVP11, and somatic hybrids (9A, 9B, 1A-25A) with RT orf125 F/R primers (see Table S5). **C.** ORF125 protein accumulation in anthers of male-sterile (SH9B) and male-fertile (SH9A) somatic hybrids. <3 and >4 indicate the size in mm of flower buds used to isolate the anthers for SH9B. For the male-fertile hybrid, a pool of flower buds with varying sizes was used. **D.** Organization of the genomic region between *orf247* and *nad4* in *tbr/*SH9B and in *cmm*/SH9A. **E.** Transmembrane domain (highlighted in violet) predicted in ORF125 with TMHMM - 2.0 webtool, spanning amino acid residues 7-24.

The *orf125* sequence spans 378 bp and is located between *orf247* and the first exon of the *nad4* gene (Fig. 2D, Fig. S1). The mitochondrial DNA and cDNA sequences of the coding region were identical, indicating no mRNA editing. Computational analysis predicted that *orf125* encodes a 125-amino-acid protein with a molecular mass of 14.66 kDa and a putative transmembrane domain between amino acids 7 and 24 (Fig. 2E, Fig. S1). Comparison of chondriome sequences from SH9A, SH9B, *cmm* and *tbr* evidenced a 4288 bp deletion within the *orf247* - *nad4* intergenic region in the male-fertile hybrid and the wild species, spanning across the *orf125* region (Fig. 2D).

### TALE-mediated knockout of orf125 induces reversion to male fertility

Two TALE-based approaches, namely TALEN and TALECD, were employed to induce mutations in *orf125* of SH9B. Mutations were induced at two different sites using each approach, resulting in a total of four independent combinations. A comprehensive analysis of the induced mutations is documented elsewhere (Nicolia et al., 2024).

At the vegetative stage, edited plants did not exhibit discernible differences from either male-sterile SH9B or male-fertile SH9A clones. On the other hand, they showed the recovery of fertility (Fig. 3). Among 17 plants transformed with TALEN sequences, four were identified as homoplasmic, with deletions of different sizes in *orf125*. Notably, all four homoplasmic plants displayed fully male-fertile phenotype. Additionally, seven heteroplasmic plants were observed, with some displaying male fertility and others male sterility (Table 1 and Table S3). All unedited regenerated plants remained male-sterile, similar to SH9B.

**Figure 3.**
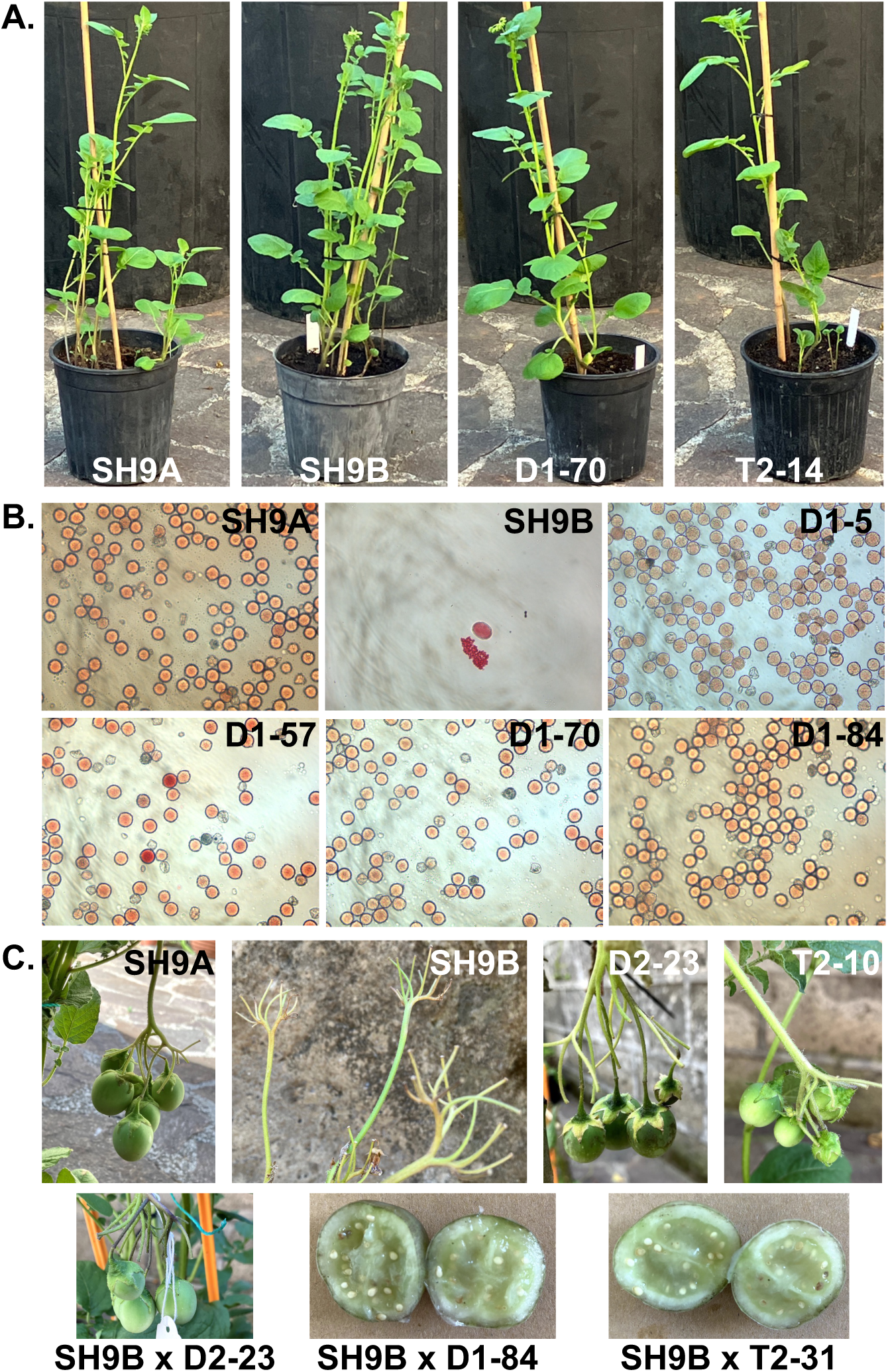
Phenotypes of edited and control plants. **A.** Tuber-derived plants grown in pots. **B.** Pollen stained with acetocarmine. **C.** Berries and seeds resulting from spontaneous selfing or crossing. D- and T-denote plants edited using the mitoTALECD or mitoTALEN approach, respectively; SH9B, male-sterile hybrid used in the editing experiments; SH9A, male-fertile hybrid.

**Table 1.**
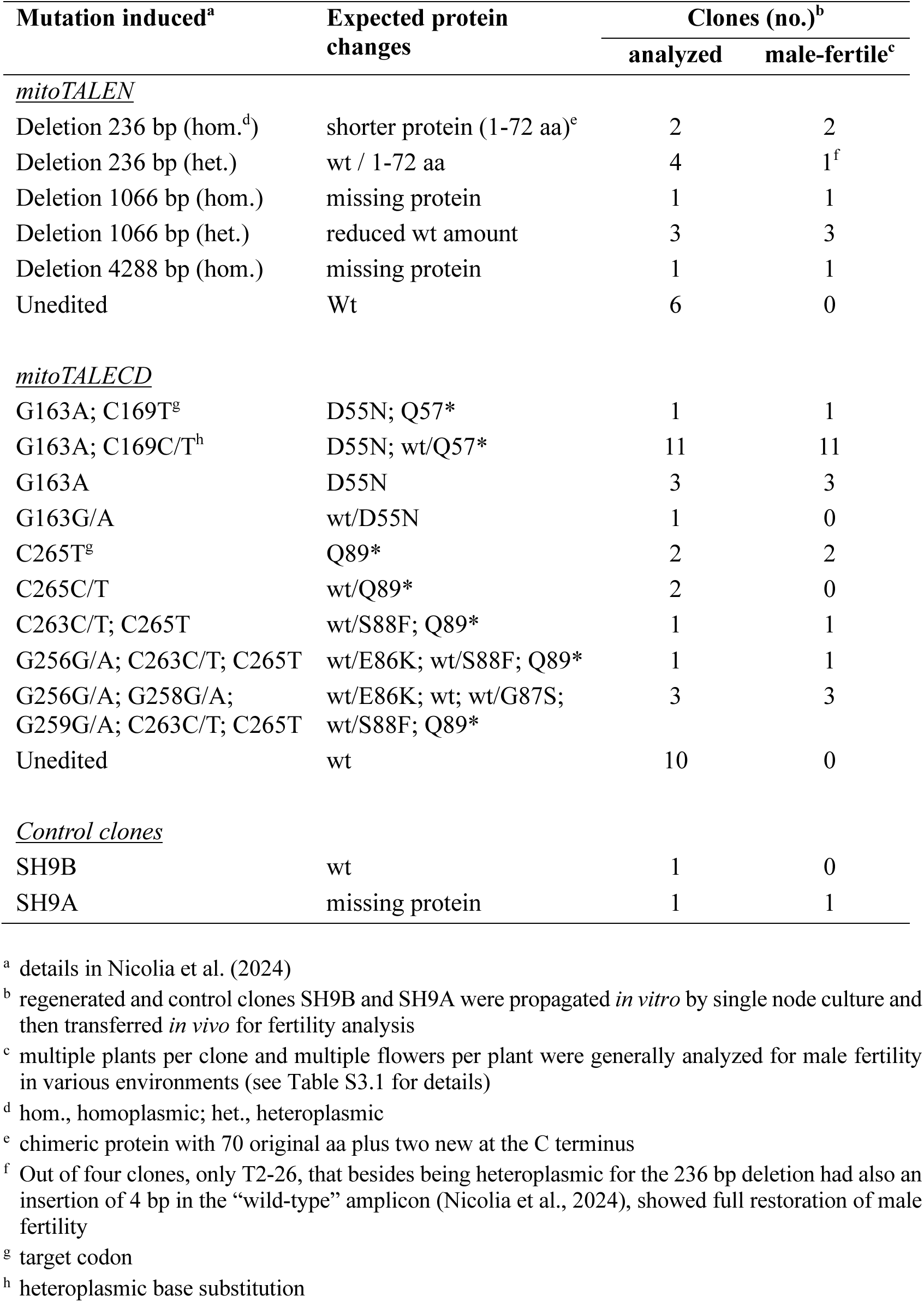
Male fertility in plants edited by mitoTALEN or mitoTALECD approach. The type of mutations induced and their homoplasmy/heteroplasmy status is also indicated.

In the TALECD experiments, 25 edited and 10 unedited plants were assessed for male fertility (Table 1 and Table S3). One plant with a homoplasmic C169T mutation and seven plants with a C265T mutation in *orf125*, resulting in a premature stop codon, demonstrated full male fertility. Intriguingly, 15 plants with a homoplasmic G163A missense mutation, leading to the D55N substitution, also exhibited male fertility. Additionally, one male-fertile plant (D1-84) displayed both the G163A and C169T mutations. However, no reversion to male fertility was observed in plants with heteroplasmic mutations or among the 10 unedited regenerated plants.

Plants derived from tuber propagation consistently maintained the phenotype observed in the previous generation (Table S3).

### Allotopic expression of orf125 in the male-fertile hybrid induces male sterility

To elucidate the role of *orf125* in inducing cytoplasmic male sterility in SH9A and explore its potential for CMS induction when expressed allotopically, we used three transformation vectors with different tissue-specific promoters and a mitochondrial signal peptide sequence. These vectors were designed for expression in photosynthetic tissues (pNS73) anther/pollen (pNS76) and tapetum (pNS79) (Fig. S2).

Independent positive transgenic plants were successfully identified via PCR for each construct (Fig. S3) and assessed for pollen production and stainability. Despite considerable variability among transgenic plants, both traits were reduced compared to SH9A (Fig. S4). SH9A produced 4.6 mg ± 0.5 of pollen per flower with 100% stainability, whereas SH9B produced very few abnormal structures that were still stainable. Allotopic expression of *orf125* in reproductive tissues significantly impacted pollen production, with reductions of up to 0.2 ± 0.1 mg and 0.5 ± 0.3 mg of pollen per flower in some NS76 and NS79 plants, respectively (Fig. S4, Fig. 4A). Pollen stainability in transgenic plants ranged from 33 to 74% (Fig. S4).

**Figure 4.**
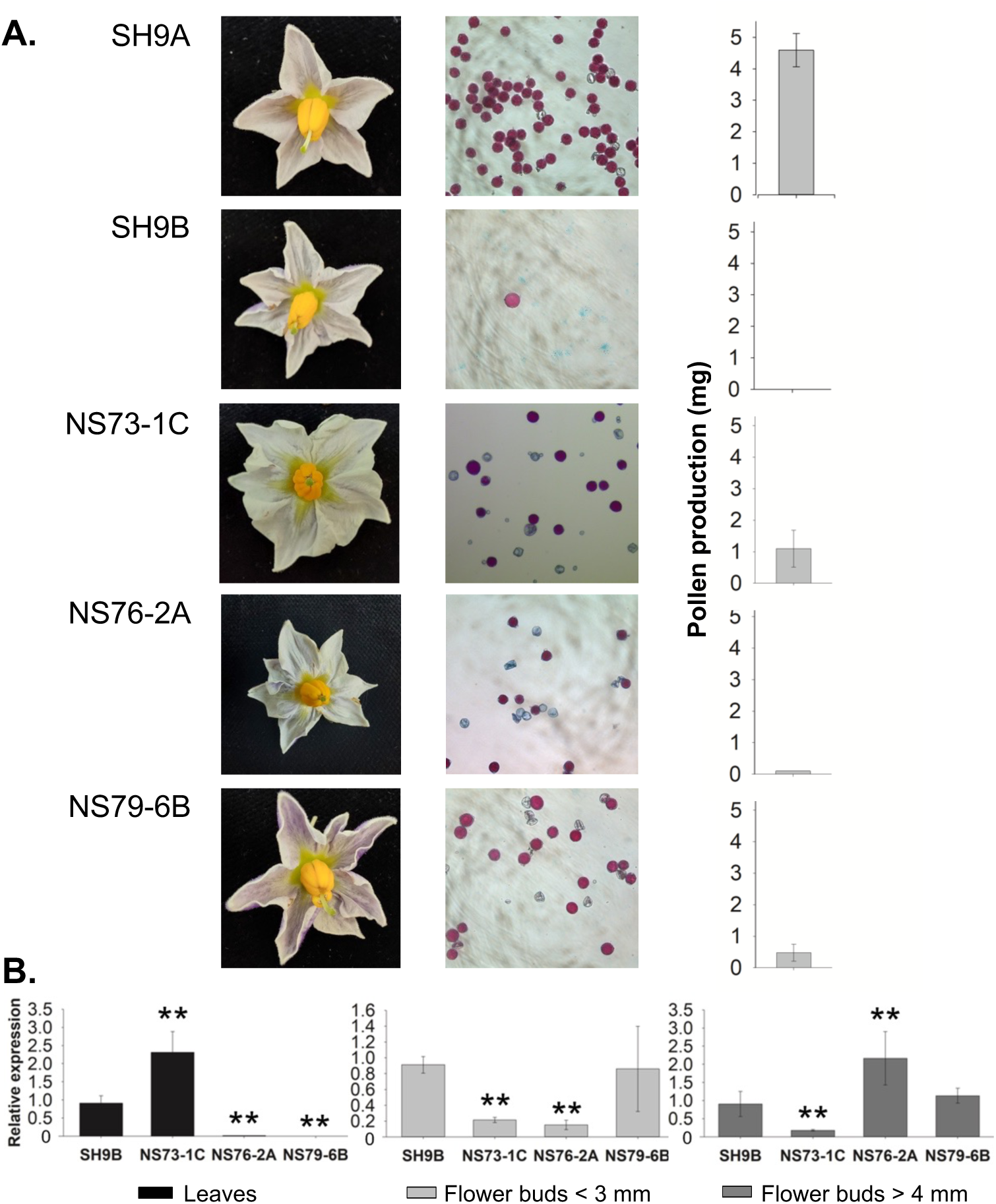
Fertility phenotype in transgenic plants overexpressing *orf125*. **A.** Flower morphology, pollen stainability and production (mg/flower) in control SH9A and SH9B somatic hybrids, and in a sample of SH9A transgenic plants expressing allotopically *orf125* under the control of constitutive (P*rbcS*, NS73), anther/pollen specific (P*lat52*, NS76) and tapetum specific (P*ta29*, NS79) promoters. **B.** *orf125* expression was determined by qRT-PCR in different tissues of SH9B and selected transgenic lines obtained with the three vectors (NS73-1C, NS76-2A and NS79-6B). The expression level was normalized using the *ef-1α* gene as reference and standardized to SH9B. *, **, P < 0.05 and P <0.01 significant differences with respect to SH9B, respectively.

As expected, *orf125* expression varied across tissues, depending on the regulatory sequence used (Fig. 4B). When controlled by the *rbcS* promoter (plant NS73-1C), *orf125* was expressed at higher levels in leaves compared to flower buds, exceeding levels observed in SH9B. By contrast, NS76-2A and NS79-6B plants showed the highest expression in flower buds. In NS76 plants, peak expression in flower buds > 4 mm, indicative of late-stage pollen, surpassed that in the male-sterile hybrid SH9B. Quantitative real-time reverse-transcription analysis demonstrated that with the tapetum specific promoter (NS79 plants), expression levels in flower buds < 3 mm were comparable to those in SH9B.

No discernible impact on pollen stainability or production was observed in control SH9A transgenic plants expressing the *gusA* marker gene under the *rbcS* or *lat52* promoters. Increased GUS expression was evident in the leaves of EF64 plants, due to the *rbcS* promoter (Fig. S5).

### The orf247-nad4 mitochondrial region exhibits distinct organizational patterns across potatoes and other Solanaceae species

The 5101 bp mitochondrial region containing *orf125*, spanning from *orf247* to the first exon of *nad4* in *tbr cvs.* Désirée and Cicero (Varré et al., 2019), was examined in other Solanaceae species via similarity search analysis (Fig. S6). Comparative analysis with *cmm* revealed that this region can be segmented into three subregions: A) nucleotides 1 to 590, B) nucleotides 591 to 4878 and C) nucleotides 4879 to the end (Fig. 5A). The B fragment was absent in *cmm*, whereas the A and C fragments were identical in both species. Analysis of sixteen *tbr* sequences from GenBank, representing 11 distinct clones, showed that these sequences were identical to the query sequence (SH9B), with only minor differences in one instance (cv. Castle Russet, MZ030732.1). The same *tbr* organization was also found in one *S. chacoense* accession (PP826245.1) (Fig. S6, Group A1).

**Figure 5.**
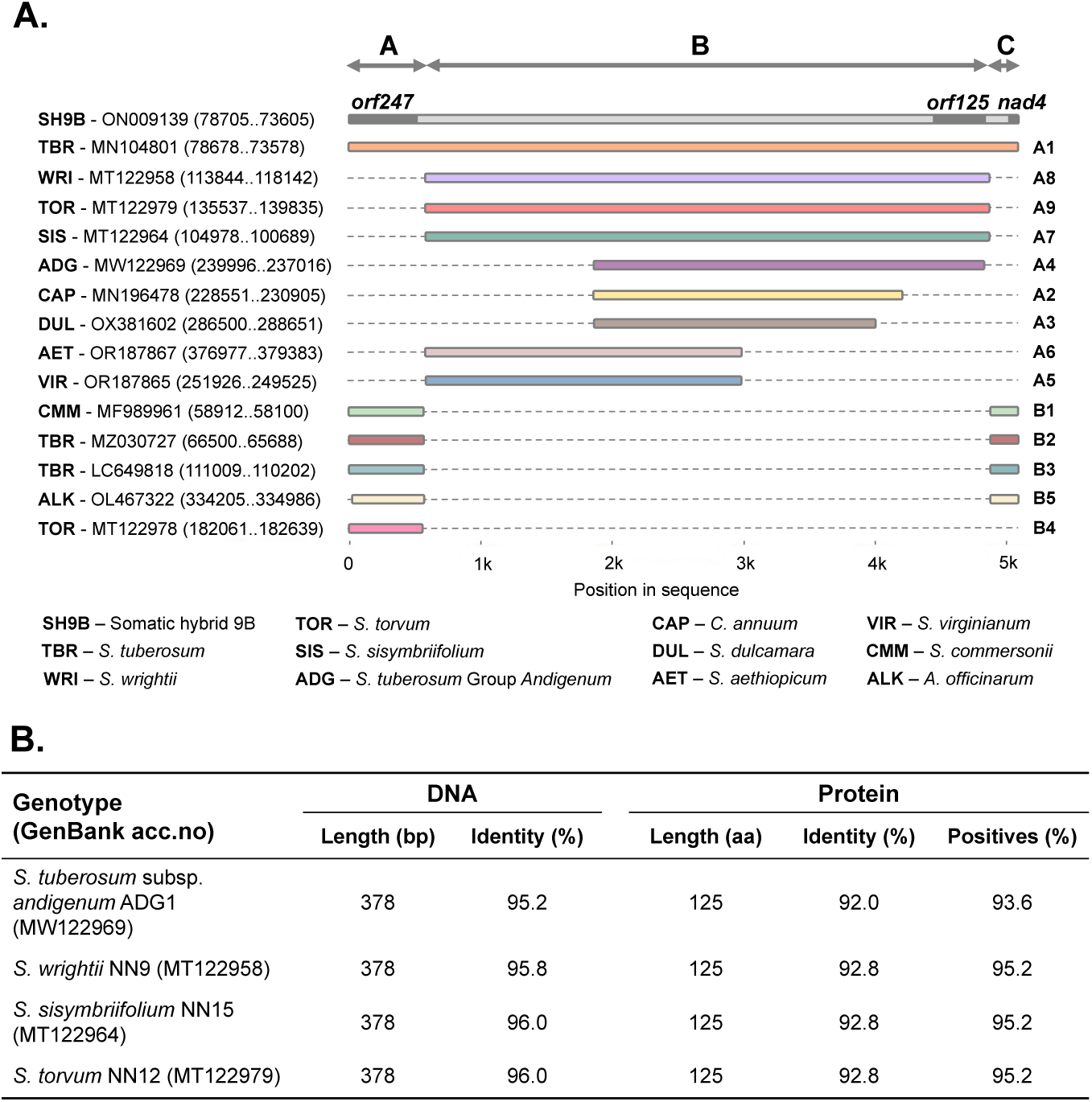
Variation in the *orf247-nad4* region among Solanaceae species. **A.** The structure of the *orf247-nad4* region is depicted in representative species groups identified through BLAST analysis, using the SH9B *orf247-nad4* sequence (P11-P3 primers, see Table S5) as the query, and Neighbor Joining (see Fig. S6). The position of *orf125* in SH9B is also indicated. A and C indicate conserved regions between *S. tuberosum* and *S. commersonii*, while B indicates the sequence present in the former but absent in the latter. **B.** Main characteristics of *orf125* nucleotide and amino acid sequences in the reported accessions of *S. tuberosum* Group *Andigenum, S. wrightii, S. sisymbriifolium* and *S. torvum*, compared to SH9B. Eleven *S. tuberosum* and one *S. chacoense* clones displayed the same *orf125* sequence as SH9B (See Fig. S6).

In contrast, several potato clones - including some from *tbr*, an accession from Group *Andigenum* (*adg2*) and others from different tuber-bearing species - cultivated and wild tomato species, pepino (*S. muricatum*), and bladder cherry (*Alkekengi officinarum* **=** *Physalis alkekengi*), all exhibited the same *orf247*-*nad4* organization as *cmm*. This configuration features contiguous A and C fragments with no B fragment present (Fig. S6, Group B). In some instances, this genomic configuration appeared duplicated in two separate regions of the chondriome. On the other hand, cultivated and wild eggplants, the ornamental *S. wrightii*, all falling under the *Leptostemonum* subgenus of the *Solanum* genus, pepper (*Capsicum annuum*), *S. dulcamara* and another accession of Group *Andigenum* (*adg1*) showed shorter B fragments (ranging from 2152 bp in *S. dulcamara* to 4299 in *S. wrightii* and *S. torvum*) (Fig. 5A). However, unlike *tbr* within the A1 subgroup, the A and C fragments in these species are positioned differently in the genome, either adjacent to each other or separated (Fig. S7).

Additional sequences, ranging from approximately 30 to 100 bp and exhibiting over 90% similarity to portions of the 5101 bp fragment, have been identified scattered across all investigated mitochondrial genomes. Furthermore, partial homologous sequences (with query coverage of 17-54% and identity exceeding 90%) have been detected on various nuclear chromosomes of *S. dulcamara*, *S. tuberosum*, *S. verrucosum*, *S. lycopersicum* and *S. pennellii*. *orf125*, however, was found as a single copy in only a few mitochondrial genomes, showing identical sequences across all twelve genotypes in group A.1. Conversely, a full-length sequence, albeit with lower identity, was found in *S. wrightii*, *S. torvum*, *S. sisymbriifolium* and the *adg1* accession (Fig. 5B). A PCR analysis carried out on a pool of tuber-bearing species (in addition to *S. etuberosum* and *S. nigrum*) provided evidence that *orf125* was successfully amplified in only two species, *S. berthaultii* (accession code *ber3*) and *S. tarijense* (*tar2*), mirroring the amplification pattern seen in SH9B and *tbr* (Désirée and SVP11). Consistent with results from *cmm* and SH9A, *orf125* was not amplified in any other species, including the *chc* accession PI 320282, which differed from the GenBank reference used for chondriome sequencing (Fig. 6A).

**Figure 6.**
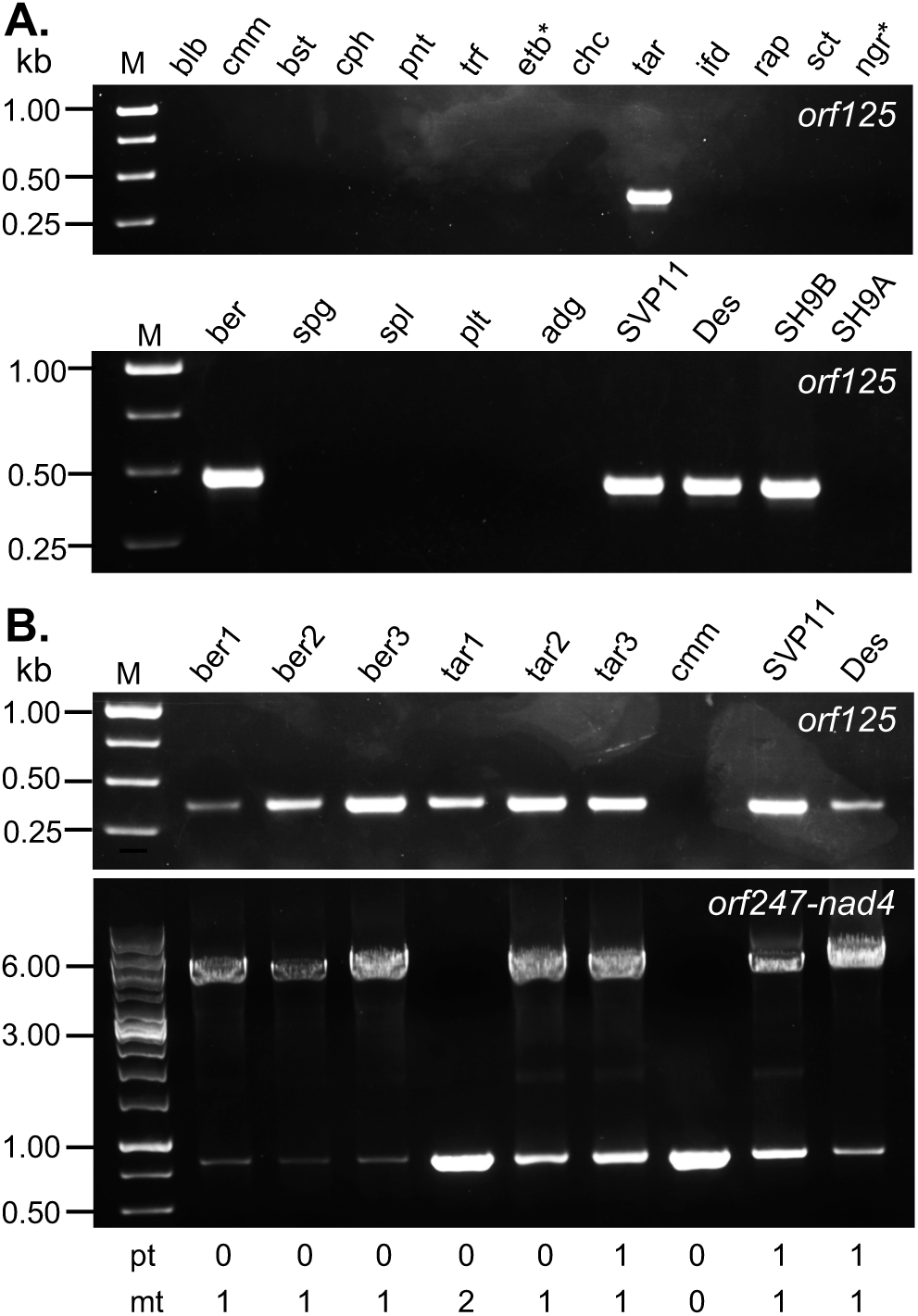
Results of PCR analyses targeting *orf125* (with primers P4-P5, see Table S5) and *orf247-nad4* (with primers P11-P3) in some tuber-bearing and other *Solanum* species (* = non-tuber-bearing). **A.** blb, *S. bulbocastanum*; cmm, *S. commersonii*; bst, S. *brachistotrichum*; cph, S. *cardiophyllum*; pnt, *S. pinnatisectum*; trf, *S. trifidum*; etb, *S. etuberosum*; chc, *S. chacoense*; tar, *S. tarijense*; ifd, *S. infundibuliforme*; rap, *S. raphanifolium*; sct, *S. sanctae-rosae*; ngr, *S. nigrum*; ber, S. *berthaultii*; spg, *S. spegazzinii*; spl, *S. sparsipilum*; plt, *S. polytrichon*; adg, *S. tuberosum* Group *Andigenum*; SVP11, Des, *S. tuberosum* Group *Tuberosum*, SH9B, SH9A, male-sterile and male-fertile somatic hybrids, respectively. **B.** ber1-3 and tar1-3 denote distinct accessions of *S. berthaultii and S. tarijense*, respectively. The designation “pt 0/1” refers, respectively, to the absence/presence of the 241 bp deletion in the plastidial genome indicative of the T-type (Hosaka, 2003), while “mt 0/1/2” indicates γ/β/a mitochondrial genome types (Scotti et al., 2007). Species names follow the taxonomy outlined by Hawkes (1994).

Based on these findings, additional amplifications of both *orf125* and the full *orf247-nad4* region were performed in two more accessions each of *ber* and *tar* (Fig. 6B). Interestingly, while all three *ber* accessions confirmed the *tbr* mitochondrial configuration (resulting in a 5101 bp long amplicon), a similar result was observed in *tar2* and *tar3* but not in *tar1*. Unexpectedly, *orf125* was amplified in all six accessions, suggesting that, *orf125* in *tar1*, unlike in *tbr* and the other accessions of *ber*/*tar*, is likely present in an alternative genomic context. Nevertheless, the *orf125* sequences from the six *ber* and *tar* accessions were identical to that of common potato.

### Wild-type and mutant ORF125 proteins show differences in overall structure

To understand the structure of ORF125 and speculate on its way of functioning, we modeled the structures of wild-type protein from *tbr* (= SH9B), its D55N mutant generated by mitoTALECD mutagenesis, and the ORF125 proteins from *adg1* and *S. wrightii/sisymbriifolium/torvum* (*wri/sis/tor*), the latter three being identical.

The protein models of wild-type from *tbr*, obtained using four independent predictors, differ in their overall architecture (see Fig. S8A-D). However, they share common features, confirming that ORF125 has a globular all-alpha structure. The initial portion of the sequence may either lack a defined conformation or function as a trans-membrane region, supporting the findings reported above.

The protein sequences from *adg1* and *wri/sis/tor* show very few amino acid differences compared to ORF125 from *tbr* (Fig. S9). While all structures are predominantly characterized by alpha helices, including an extended helix on the N-terminal side, these minor amino acid differences might affect the relative arrangement of secondary structure domains (Fig. S8E-H). The superimposition of wild-type and D55N mutant reveals a high Root Mean Square Deviation (RMSD) of 10.01 Å. Similarly, superimposing *tbr* ORF125 with those from *wri/sis/tor* or *adg1* reveals high RMSD values (9.24 Å and 8.18 Å respectively). The structures of ORF125 from *adg1* and *wri/sis/tor* are more similar, with a RMSD of 3.77 Å. However, when superimposition is performed by excluding either the N- or the C-terminal regions (amino acids 1-41 or 42-125, respectively) the RMSD values between wild-type and mutant forms decrease (Fig. S10). This suggests that the high RMSD values observed in the full superimposition are due to rearrangements in these terminal regions.

## Discussion

### orf125 as mitochondrial causal determinant of male sterility in tbr (+) cmm hybrids

The evaluation of the selective presence and expression of candidate *orfs* in male-sterile/fertile hybrids, along with parental species, strongly suggested that *orf125* could be the key mitochondrial determinant of male sterility in interspecific *tbr* (+) *cmm* hybrids. This was unequivocally confirmed through mitochondrial genome editing techniques (mitoTALEN and mitoTALECD) and allotopic expression transgenic approaches.

SH9B plants edited by mitoTALEN exhibited homoplasmic deletions of varying lengths within the target region containing *orf125*. Of particular interest, one mitoTALEN-edited plant (T2-12) displayed the same 4288 bp deletion that differentiates *cmm* and SH9A from *tbr* and SH9B. Conversely, base editing techniques produced a range of SH9B plants with precise missense or nonsense mutations in *orf125* (Nicolia et al., 2024). Recent advancements in mitoTALEN technology have shown its effectiveness in various plant species, enabling the targeted inactivation of essential respiratory chain mitochondrial genes and *orfs* potentially associated with CMS (Kazama et al., 2019; Arimura et al., 2020; Kuwabara et al., 2022; Ayabe et al., 2023; Forner et al., 2023; Xu et al., 2024). On the other hand, organellar base editing using TALE-DddA fusion proteins has primarily targeted plant plastomes (reviewed by Nakazato and Arimura, 2024) with limited data available on mitochondrially edited plants (Nakazato et al., 2022). Interestingly, not only did the potato homoplasmic plants with physical deletions or induced premature stop codons in *orf125* revert to male fertility, but also those with a single amino acid substitution (aspartic acid to asparagine) at position 55. This substitution likely alters the structure of the ORF125 protein, impacting its function. Specifically, the structural modification appears to affect the orientation between the long N-terminal helix and the C-terminal domain, as indicated by the reduced RMSD values when comparing structures with either the N- or the C-terminal regions excluded. The relative rearrangement may affect ORF125 ability to interact with other proteins, leading to varied functional behaviors across different species and in its mutated form.

The putative role of *orf125* during meiosis was substantiated by the significant reduction in pollen production, rather its stainability, when expressed in the nucleus and as a mitochondrial-targeted protein in anther cells of the transgenic SH9A somatic hybrid. This mirrors the phenotype seen in male-sterile somatic hybrids, where pollen or pollen-like structures were scarce but partially stainable (Conicella et al., 1997). As in similar cases (Hanson and Bentolila, 2004), however, complete sterility was not achieved, probably due to differences in the expression of the transgenic protein allotopically expressed in the nucleus and targeted to mitochondria, compared to the native mitochondrial ORF125.

The same *orf125* sequence found in male-sterile *cmm* (+) *tbr* somatic hybrids was also identified in several *4x* and 2*x S. tuberosum* clones, which are generally male-fertile, as evidenced by their use as either male or female parents in breeding schemes (*e.g. cvs.* Désirée, Spunta and Atlantic, https://www.plantbreeding.wur.nl/PotatoPedigree/index.html). This suggests that *orf125* is not sufficient to induce male sterility; interaction with unidentified nuclear genes present in some species may be necessary to induce male sterility in hybrid genotypes. Structural investigations of ORF125 somehow support this, as the reference proteins selected by different predictors were significantly larger, containing functional domains similar to ORF125, indicating potential interactions or modifications that could affect sterility. The functions of such proteins or their ORF125-like domains typically involve interactions with other molecules, such as protein-protein interactions, nucleic acid binding, or roles as transport and trafficking. Variations in protein folding observed in different forms of ORF125 could significantly impact these interactions and functions.

### Evolutionary insights

Patterns of variation in the organization of the *orf247-nad4* region have been observed not only within potato species, but also across other Solanaceae family members. The intergenic sequence found in common potato appears to have emerged multiple times throughout evolution, with varying length and content, likely due to recombination events involving homologous sequences scattered throughout the chondriome. In tuber-bearing *Solanum* species, a conserved organization of this genomic region was found in tetraploid and diploid *tbr* clones with T cp-type and β mt-type (except for cv. Russet Castle which has a W/γ cytoplasm (Hoopes et al., 2022), as well as in one *chc* accession and several *ber*/*tar* accessions. These clones also showed the presence of the same *orf125* sequence. By contrast, species and clones with variant cytoplasms, as determined through molecular and/or pedigree analyses, exhibited an alternative genomic organization, lacked *orf125*, and grouped separately (Lössl et al., 1999; Scotti et al., 2007; Gargano et al., 2012; Cho et al., 2016; Cho et al., 2017; Cho et al., 2018; Varré et al., 2019; Achakkagari et al., 2020; Achakkagari et al., 2021a; Achakkagari et al., 2021b; Hoopes et al., 2022; Sanetomo et al., 2022).

One out of two *Andigenum* accessions available in GenBank displayed a variant form of *orf125* and an alternative local genomic organization. However, the two accessions have been shown to differ not only in chloroplast type but also in mitochondrial genome sequence (Achakkagari et al., 2020; Achakkagari et al., 2021b) highlighting the large cytoplasmic variability known in Andean tetraploid potatoes (Hosaka and Hanneman, 1988; Gavrilenko et al., 2013). Similarly, only one of the two *chc* accessions, either present in GenBank or analyzed in this study, carried the *orf125* sequence. Despite having the same *orf247-nad4* region organization as *tbr*, both plastidial and mitochondrial genomes of *chc* were found to be divergent from *tbr* (Scotti et al., 2007; Kim and Park, 2019). Hence, it is unlikely that *orf125* in *tbr* originated from *S. chacoense*.

By contrast, the *S. berthaultii* complex, which includes both *S. berthaultii* and *S. tarijense,* now recognized as a single species (Spooner et al., 2007), is a more likely candidate. This complex, in fact, exhibited the “Tuberosum”-type plastid DNA in approximately 18% of the accessions investigated (Spooner et al., 2014) and has a plastome sequence close to that of *tbr* (Kim and Park, 2019).

Among the six accessions analyzed in this study, *tar3* (PI442689) showed the 241 bp deletion indicative of the T chloroplast-type (Hosaka, 2003), while *tar1* (PI265577) featured the variant mitochondrial genome α instead of the β-type observed in the other five accessions (Lössl et al., 2000; Scotti et al., 2007). The presence of *orf125* in all six *ber/tar* accessions suggests that the emergence of *orf125* either predates the plastid differentiation in *S. berthaultii/tarijense* spanning central Bolivia to northwest Argentina or results from repeated independent recombination events. Further studies involving a broader range of accessions from the *S. berthaultii* complex are therefore warranted.

Tetraploid accessions of *S. tuberosum* Group *Andigenum* displayed high diversity in plastid genomes with predominance of the A-type, whereas *S. tuberosum* Group *Chilotanum* predominantly featured almost exclusively the T-type (Hosaka and Hanneman, 1988) (Fig. S11). Among wild species, the T-type was only found in *S. neorossi* and *S. berthaultii/tarijense,* leading to the hypothesis that the Chilean potato originated from hybridization between 4*x S. tuberosum* Group *Andigenum* and a 2*n* egg-producing clone of *S. berthaultii/tarijense*, which served as the cytoplasm donor (Grun, 1990; Hosaka, 2003; Hosaka and Sanetomo, 2009; Gavrilenko et al., 2013; Spooner et al., 2014). The presence of *orf125* in *ber/tar*, particularly in *tar3* in combination with other cytoplasmic markers specific to *tbr*, largely supports this hypothesis and provides insights into the origin of mitochondrial factors contributing to genic-cytoplasmic male sterility observed in crosses between *tbr* and *adg* (or some wild species) (Hermundstad and Peloquin, 1985; Grun, 1990; Anisimova and Gavrilenko, 2017; Goryunova et al., 2023).

A plausible scenario suggests that initial male sterility, stemming from the interplay between *orf125* from *berthaultii* and nuclear genes from the *Andigenum* parent, combined with vegetative propagation of superior (heterotic) genotypes, facilitated the emergence and isolation of the new species. This led to the widespread prevalence of the T/β cytoplasm, first in *S. tuberosum* Group *Chilotanum* and later in *S. tuberosum* Group *Tuberosum* (Fig. S11). This bottleneck effect may have been further augmented by the beneficial impact of the T/β cytoplasm on some agronomic traits (Hosaka et al., 2018; Goryunova et al., 2023). Based on the observed increase in male fertility of Tuberosum - Neo-Tuberosum progenies compared to Tuberosum - Andigena progenies after selection for tuberization under long days, which suggests pleiotropic or linkage effects between genes controlling photoperiod response and male fertility (Vilaró et al., 1989), it can be hypothesized that also the recovery of a certain degree of male fertility in chilean landraces likely occurred later through natural and human selection that favored tuberization under long-day conditions typical of southern South America.

The “Tuberosum”-type nuclear-cytoplasmic male sterility induced by the T/β cytoplasm was later incorporated into European potato cultivars and those developed globally following the introduction of Chilean potato to Europe. However, its prevalence now varies, influenced by the use of other species as cytoplasm donors for transferring disease resistance genes and overcoming male sterility (Hosaka and Sanetomo, 2012).

Further investigations are, however, necessary to further elucidate functional mechanisms of CMS induction by *orf125*, its distribution in tuber-bearing *Solanums* and contribution to the evolution of common potato.

## Materials and Methods

### Plant material

Genetic materials used in this study are listed in Table S4. *Solanum* species and clones were provided by Dr. J.B. Bamberg (Potato Introduction Station, Sturgeon Bay, Wisconsin) and Dr. G. Ramsay (Commonwealth Potato Collection, Dundee, UK). Somatic hybrids SH9A (male-fertile), SH1A, SH7A, SH9B, SH12A and SH25A (all male sterile) were obtained via protoplast fusion between *tbr* dihaploid clone SVP11 and *cmm* accession PI 243503 (Cardi et al., 1993). SH9A and SH9B originated from the same callus.

### Isolation of mitochondria and mtDNA extraction

Potato mitochondria were isolated from tubers using a juice extractor, followed by homogenization in 3x grinding buffer pH 7.5 (0.9 M sucrose, 90 mM sodium pyrophosphate, 6 mM EDTA, 12 mM cysteine, 15 mM glycine, 2% PVPP 360,000, 0.9% BSA, 6 mM β- mercaptoethanol). After washing and differential centrifugation steps, mitochondria were purified on a discontinuous Percoll gradient (14-28-45% v/v) at the 28-45% interface. Finally, a DNase I (0.5 mg/100 g tubers; Invitrogen, USA) treatment was performed for 45-60 min at 37 °C (Varré et al., 2019). Mitochondrial DNA extraction followed the method described by Scotti et al., (2001). Briefly, pellets were resuspended in 1 ml lysis buffer (25 mM Tris-HCl pH 8.0, 20 mM EDTA pH 8.0, 0.5% SDS) containing Proteinase K (50 µg/ml) and RNase (25 µg/ml), then incubated for 1 h at 37 °C. Subsequently, 0.1 volume of 2 M ammonium acetate was added, and nucleic acids were extracted with an equal volume of TE-saturated phenol/chloroform (50:50) and centrifuged at 10,000xg for 10 min at 10 °C. Finally, nucleic acids were precipitated by adding 2 volumes of 100% ethanol and overnight incubation at -20 °C. After centrifugation at 16,000x*g* for 15 min, the pellet was re-dissolved in water.

### Mitogenome assembly, annotation, and identification of synteny blocks

The mtDNA isolated from SH9A and SH9B somatic hybrids was sequenced using Illumina and PacBio technologies. Illumina reads were quality checked using FastQC v0.11.9 (Andrews, 2010) and Trimmomatic v0.39 (Bolger et al., 2014). Potential contaminating sequences were screened using Mash v2.3 (Ondov et al., 2019) and mitochondrial reads were filtered out using *BBDuk* from the BBMap v38.95 package (Bushnell, 2014). PacBio reads were quality processed with Filtlong (https://github.com/rrwick/Filtlong), discarding reads shorter than 5 kbp and the worst 10% of bases. Mitochondrial DNA was assembled Unicycler v0.5.0, in “normal” mode (Wick et al., 2017), with SPAdes v3.15.4 (Prjibelski et al., 2020) for short reads and minimap and miniasm (Li, 2016) for long reads. Polishing was done with Racon v1.5.0 (Vaser et al., 2017). After assembling and polishing, mitochondrial DNA was annotated by comparison with previously annotated potato mtDNA sequences (accession numbers: MN104801, MN104802, MN104803), and synteny blocks identified using Sybelia v3.0.7 (Minkin et al., 2013) (min block size: 500 bp) and graphed with *RIdeogram* v0.22 (Hao et al., 2020).

### PCR, RT-PCR and qRT-PCR analyses

Genomic DNA (gDNA) extraction was performed using the DNeasy Plant Mini kit (Qiagen, Germany) according to the manufacturer’s instructions. and used for PCR analyses. These were carried out using either Taq recombinant or Phire Hot Start II DNA Polymerase, depending on the target length (Invitrogen, USA).

cDNA was synthetized from 1 µg RNA treated with DNase I and used for RT-PCR and qRT-PCR analyses. RNA extraction was carried out using the RNeasy Plant Mini kit (Qiagen, Germany) following the manufacturer’s guidelines. The quality of the extracted RNA was assessed through Nanodrop (Thermo Fisher, USA) measurements, while its integrity was confirmed via agarose gel electrophoresis. To eliminate any residual DNA, a DNase I treatment (Invitrogen, USA) was performed as per manufacturer’s instructions. Subsequently, cDNA synthesis was performed for all samples, using 1 µg of RNA following the protocol outlined in the RevertAid 1^st^ strand cDNA synthesis kit (Thermo Fisher Scientific, USA). The quality of the resulting cDNA was further evaluated through RT-PCR amplification of the *18S rRNA* gene, with amplification products visualized via agarose gel electrophoresis. Diluted cDNA (1:20) was then used as a template for quantitative real-time reverse-transcription PCR, using the Platinum™ SYBR™ Green qPCR SuperMix-UDG (Applied Biosystems, USA).

All reactions were prepared using 6.25 µl of 2x SYBR green dye, a primer mix of 0.6 µM of forward and reverse primers, and 4.5 µl template cDNA. These reactions were run on the 7900HT Fast Real Time PCR system (Applied BioSystems, USA). The program cycle parameters were configured as follows: 50 °C for 2 min (stage 1), 95 °C for 10 min (stage 2), 95 °C for 15 sec and 60 °C for 1 min (stage 3 repeated 40 times), 95 °C for 15 sec, 60 °C for 15 sec and 95 °C for 15 sec (stage 4). Melting curve analysis was performed to assess the specificity of the reactions. Relative expression levels were determined using the 2^-ΔΔCT^ method (Livak and Schmittgen, 2001) with *ef-1α* serving as the internal control. Data representing three biological replicates and three technical replicates are presented as means. Differences in means between transgenic plants and SH9B were assessed using Student’s t-test. All statistical calculations were done using Microsoft Excel for Microsoft 365 MSO and SigmaPlot 12.0.

### Western blotting

Proteins were extracted from anthers isolated from flower buds (< 3 or > 4 mm in size) of SH9A and SH9B somatic hybrids by homogenization in 0.1 M Tris-HCl (pH 7.8) containing 0.2 M NaCl, 1 mM EDTA, 0.2% Triton X100, 2% SDS, 2% β-mercaptoethanol, 1 mM PMSF, 1x proteinase inhibitor cocktail. After centrifugation at 20,000x*g* for 15 min at 4 °C, the supernatant, containing the extracted proteins, was collected. Protein concentrations were determined using the Bradford method with Bio-Rad protein assay reagent (Bio-Rad, USA) and bovine serum albumin (BSA) as the reference standard. Thirty μg of protein samples were loaded onto an 18% polyacrylamide gel. Following protein separation, they were transferred to a nitrocellulose membrane for 55 min at 100 V and room temperature.

The membrane was blocked for 2 h at room temperature in PBS containing 3% BSA and 0.1% Tween20, then incubated overnight at 4 °C with a custom-made primary antibody (Primm, Italy) diluted 1:100 in blocking buffer. Following washing steps, the membrane was incubated with anti-rabbit secondary antibody (1:60,000) diluted in PBS containing 0.1% Tween20 and 5% skim milk for 1 h at room temperature. Chemiluminescence signals were detected using the Western Blotting ECL Prime kit (GE Healthcare, USA) and visualized with a ChemiDoc^TM^ XRS^+^ (Bio-Rad). Image analysis was performed using the Image Lab^TM^ Software (Bio-Rad).

*Construction of transformation vectors, plant transformation, mitochondrial genome editing* Vectors designed to elucidate the role of *orf125* in inducing cytoplasmic male sterility in potato contained the *orf125* coding sequence, along with the mitochondrial signal peptide sequence from the yeast *coxIV* gene, driven by three tissue-specific promoters (Fig. S2): P*rbcS* from *Chrysanthemum morifolium* for expression in photosynthetic tissues (pNS73); P*lat52* and P*ta29* from *S. lycopersicum* tailored for anther/pollen and tapetum specific expression (pNS76 and pNS79), respectively (Twell et al., 1989; Mariani et al., 1990). These vectors were used to transform the male-fertile somatic hybrid SH9A.

The *orf125* coding sequence underwent PCR amplification to incorporate *Nco*I and *Bgl*II restriction sites, followed by subcloning into the commercial intermediate plasmid ImpactVector 1.5 (Wageningen, The Netherlands, http://www.impactvector.com). This plasmid harbors the promoter, 5’-UTR, and terminator of the *rbcS* gene, along with two tags at the C-terminus (c-myc and 6xHis), and the mitochondrial signal peptide sequence derived from the yeast *coxIV* gene. Additional intermediate vectors were developed by replacing the P*rbcS* promoter region (specific for photosynthetic tissues) with the promoter region of *Solanum lycopersicum lat52* and *ta29* genes, which were amplified *via* PCR to introduce *Asc*I and *Xba*I restriction sites. The tomato P*ta29* promoter has been kindly provided by Prof. Ivo Rieu, Radboud Institute for Biological and Environmental Sciences, Radboud University, Nijmegen, Netherlands. The correct sequences of expression cassettes were verified by Sanger sequences of all intermediate vectors.

The three expression cassettes were ligated as *Asc*I-*Pac*I fragment into the binary vector pBIN plus from the commercial ImpactVector kit (Fig. S2), resulting in the generation of pNS73, pNS76 and pNS79 vectors. Subsequently, binary vectors were transferred into *Agrobacterium tumefaciens* strain LBA 4404 and used to transform explants of *in vitro*-grown male-fertile somatic hybrid SH9A, according to the protocol by Andersson et al., (2003). Control plants were produced by transforming SH9A with binary vectors derived from Impact Vectors 1.5 and containing the *gusA* gene under the control of *rbcS* (pEF64) and *lat52* (pNS78) promoters (Fig. S5).

The presence of transgenes in kanamycin resistant plants has been confirmed by primers P*rbcS* F, P*lat52* F, Pt*a29* F and orf125 *Bgl*II R landing on the 5’ promoter and 3’ coding sequences of *orf125* cloned in pNS73, pNS76 and pNS79, respectively (Table S5).

TALE sequences linked to *Fok*I (mitoTALEN) or to a DddA cytidine deaminase (mitoTALECD) were designed in two regions of the *orf125* gene to achieve, respectively, a double-strand break or a stop codon by targeted base editing. Sequences for editing were cloned in plant expression vectors containing the 35S promoter and the N-terminal pre-sequence of the Arabidopsis mitochondrial ATPase delta-prime subunit. Production and molecular characterization of edited plants have been described previously (Nicolia et al., 2024).

### Fertility assessment

Transgenic SH9A and edited SH9B plants were assessed for pollen production and stainability, and compared to original SH9A and SH9B somatic hybrids. Pollen was collected from plants grown in greenhouse and growth-chamber. Pollen production was estimated using 10-15 flowers from 4-5 plants per genotype. Pollen viability was evaluated by staining either with acetocarmine or Alexander method (Alexander, 1969).

### Model construction and analysis

ORF125 protein sequences were aligned using Clustal Omega (Madeira et al., 2022). Models of ORF125 from *Solanum tuberosum* Group *Tuberosum* have been performed exploiting four different bioinformatic tools: I-TASSER (Yang and Zhang, 2015), SWISS MODEL (Waterhouse et al., 2018), AlphaFold2 (ColabFold v1.5.5) (Mirdita et al., 2022) and Phyre2 (Kelley et al., 2015), simply giving the protein sequence as input. Models of ORF125 D55N mutated form, and of ORF125 proteins from *S. tuberosum* Group *Andigenum* and *S. wrightii/sisymbriifolium/torvum* have been built using only AlphaFold2. All settings have been left as default. Z-score of models obtained have been evaluated by Prosa-web (Sippl, 1993).

Structures visualization and images creation were performed using BIOVIA Discovery Studio Visualizer (Dassault Systèmes), and Chimera 1.14 (Pettersen et al., 2004).

### BLAST analyses

BLAST+ version 2.12.0 was used to compare the 5101 bp region containing *orf125* (from position 78705 to 73605 of sequence ON009139) with nucleotide sequences of the Solanaceae family available in GenBank. The parameters were set to include matches ≥ 90% identity and a minimum length of 140 bp. Entries related to somatic hybrids, CMS genotypes, and the nuclear genome were excluded. The BLAST results were visualized using Inkscape software (https://inkscape.org/). Dot plots comparing the 5101 bp region with GenBank sequences MT122958, MT122979, MT122978, MT122964, MW122969, MN196478, OR187867, OR187865, OX381602, were generated using dot_plot_like_in_BLAST.py (https://github.com/shelkmike/Dot_plot_like_in_BLAST).

### Data Availability

The two mitogenomes reported in this paper have been archived in GenBank under the following accession numbers: ON009139, ON009140, ON009141 for SH9B and ON682437, ON682438, ON682439, ON682440 for SH9A. PacBio and Illumina reads have been submitted to the Sequence Read Archive (SRA) under the project accession number PRJNA1114443. All additional data can be found within the article and/or SI Appendix.

## Funding

This study was carried out within the Agritech National Research Center and received funding from the European Union Next-GenerationEU (PIANO NAZIONALE DI RIPRESA E RESILIENZA (PNRR) – MISSIONE 4 COMPONENTE 2, INVESTIMENTO 1.4 – D.D. 1032

17/06/2022, CN00000022). This manuscript reflects only the authors’ views and opinions, neither the European Union nor the European Commission can be considered responsible for them.

## Author Contributions

TC and NS conceived and developed the original concept with contributions from RT and NDA; NS and RT designed and performed molecular biology and overexpression experiments; NDA and GA designed and performed sequencing and bioinformatic analyses; TC, SA and AN designed and performed editing experiments; AF and DG designed and performed protein modelling analyses; LS and RP propagated plants and carried out fertility analyses; TC and NS coordinated all research; TC wrote the manuscript with contributions from NS, RT, NDA, AN and AF; All authors revised and commented the manuscript.

## Acknowledgments

We thank Prof. Ivo Rieu, Radboud University, Nijmegen, Netherlands, for kindly providing the P*ta29* promoter, and Prof. D. Carputo, University of Naples Federico II, Portici, Italy, and Dr. Maria Stefania Grillo, CNR-IBBR, Portici, for kindly revising the manuscript.

