## Supplemental material for "The mitochondrial *orf125* affects male fertility of *Solanum tuberosum* (+) *S. commersonii* somatic hybrids and participates in the onset of “Tuberosum”-Type CMS and evolution of common potato"

### Supplemental information

**ATG**AATATCTTTGATATTTTCACGATTGCCTATTCAATGATTTTTTTTGTCTCTTTCACTTTTTTCGTA
GCCAGACACAAACTTTCTTCCTATTTATCCTTTAATGACCTTCTGACTAGAGAAGAATACCTTCGGCTG
CTTGTCCCGGAGGCCCTTCAGAAGGATATTCAACTAATACTAAACTTATATTTGACTAACTTGGGCGAG
AGGATGCCCCAAGGGCACTCTTGGCAAGAGCTCGCCCACAAAATTCACGAGGGCTCTCAAGACAAGGAG
TTCCTTCTTTCCATTATATCTGACATTGTTTCTAATGGAATGTCAAGTGAATGGGTCTTGATGGCTTTG
CAAATCCTGAAAGATTGGACCTTTCCTCTA**TAA**

Predicted transmembrane domain

MNIFDIFTIAYSMIFFVSFTFFVARHKLSSYLSFNDLLTREEYLRLLVPEALQKDIQLILNLYLTN
LGERMPQGHSWQELAHKIHEGSQDKEFLLSIISDIVSNGMSSEWVLMALQILKDWTFPL

**Figure S1.** Nucleotide sequence of *orf125* and deduced protein sequence in SH9B. Start and stop codons are indicated in bold, while the dotted line indicates the predicted transmembrane domain.

**A.**

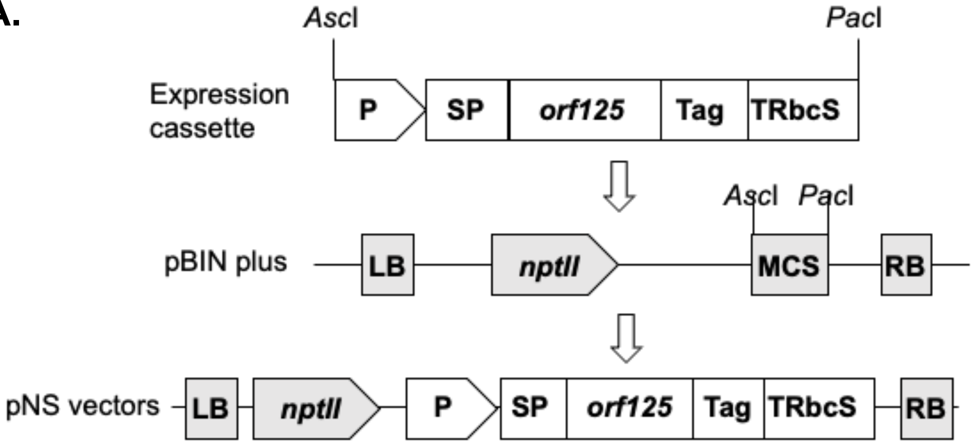


**B.**

| Vector | Promoter[a] | Signal peptide[b] | Transgene | Tag | Terminator[c] |
|---|---|---|---|---|---|
| pNS73 | P*rbcS* | mt CoxIV | *orf125* | c-myc /6xHis | T*rbcS* |
| pNS76 | P*lat52* | mt CoxIV | *orf125* | c-myc /6xHis | T*rbcS* |
| pNS79 | P*ta29* | mt CoxIV | *orf125* | c-myc /6xHis | T*rbcS* |

[a] Promoter region of *Chrysanthemum morifolium* small subunit of Rubisco (photosynthetic tissues specific) or promoter region of *Solanum lycopersicum lat52* gene (anther-pollen specific) or *ta29* gene (tapetum-specific)

[b] Signal peptide from yeast mitochondrial CoxIV protein

[c] Terminator region of *Chrysanthemum morifolium* small subunit of Rubisco

**Figure S2.** Schematic representation (**A.**) and description (**B.**) of vectors used for overexpression of *orf125* CMS-candidate gene in the male-fertile somatic hybrid SH9A.

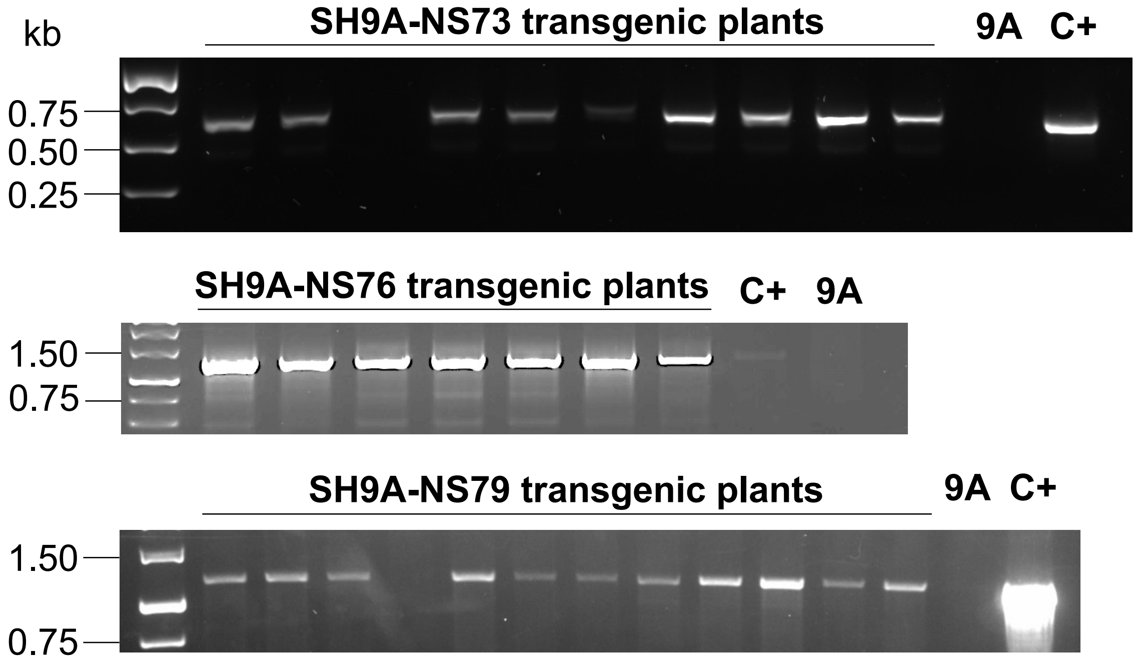


**Figure S3.** Selection of positive *orf125* transgenic plants after transformation with pNS73, pNS76 and pNS79 vectors. The transgene fragment was amplified with P*rbcS* F/*orf125 Bgl*II R , P*lat52* F/*orf125 Bgl*II R and P*ta29* F/*orf125 Bgl*II R primers, respectively (see Table S5). 9A, SH9A; C+, DNA vectors.

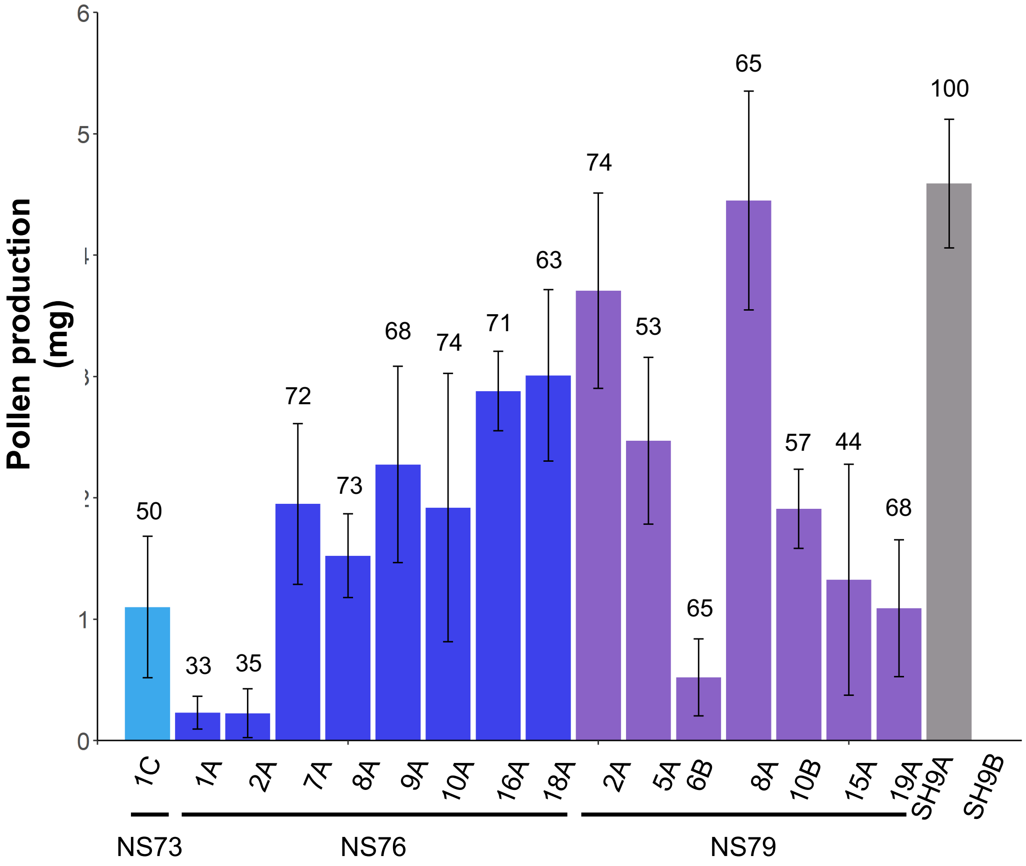


**Figure S4.** Male fertility of SH9A transgenic plants expressing *orf125* under the control of P*rbcS* (NS73), P*lat52* (NS76) and P*ta29* (NS79) promoters. Bars represent pollen production (mg/flower ± SD). Numbers above bars indicate average pollen stainability (%).

**A.**

| Vector | Promoter | Signal peptide | Transgene | Tag | Terminator |
|---|---|---|---|---|---|
| pEF64 | P*rbcS* | mt CoxIV | *gus*-int | c-myc /6xHis | T*rbcS* |
| pNS78 | P*lat52* | mt CoxIV | *gus*-int | c-myc /6xHis | T*rbcS* |

**B.**

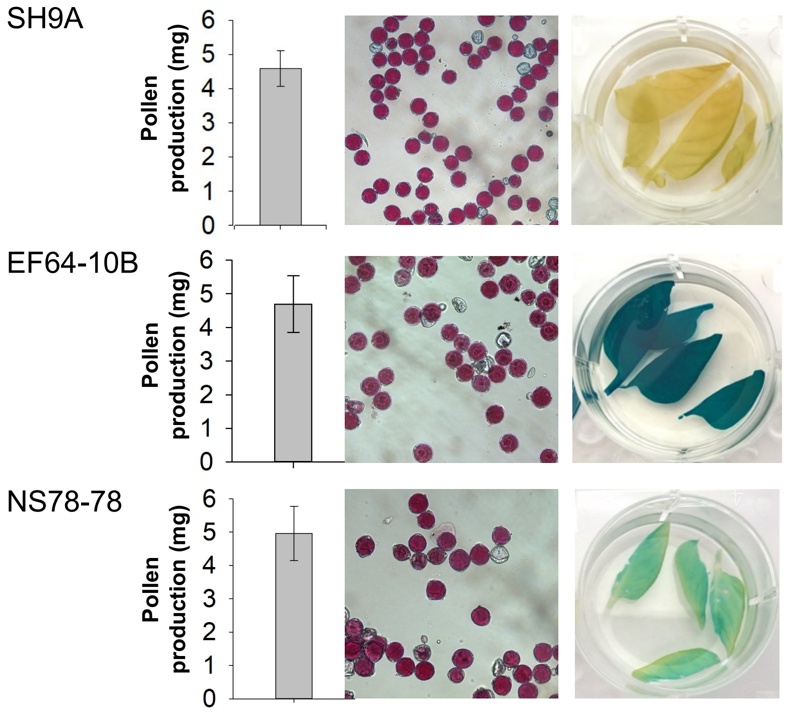


**Figure S5.** Effects of the expression of *gus* gene on pollen production and stainability. A. Description of vectors used for overexpression of the *gus* gene. B. Pollen production and stainability, and results of histochemical GUS assay in leaves of the male-fertile somatic hybrid SH9A and of selected transgenic plants (EF64-10B and NS78-78).

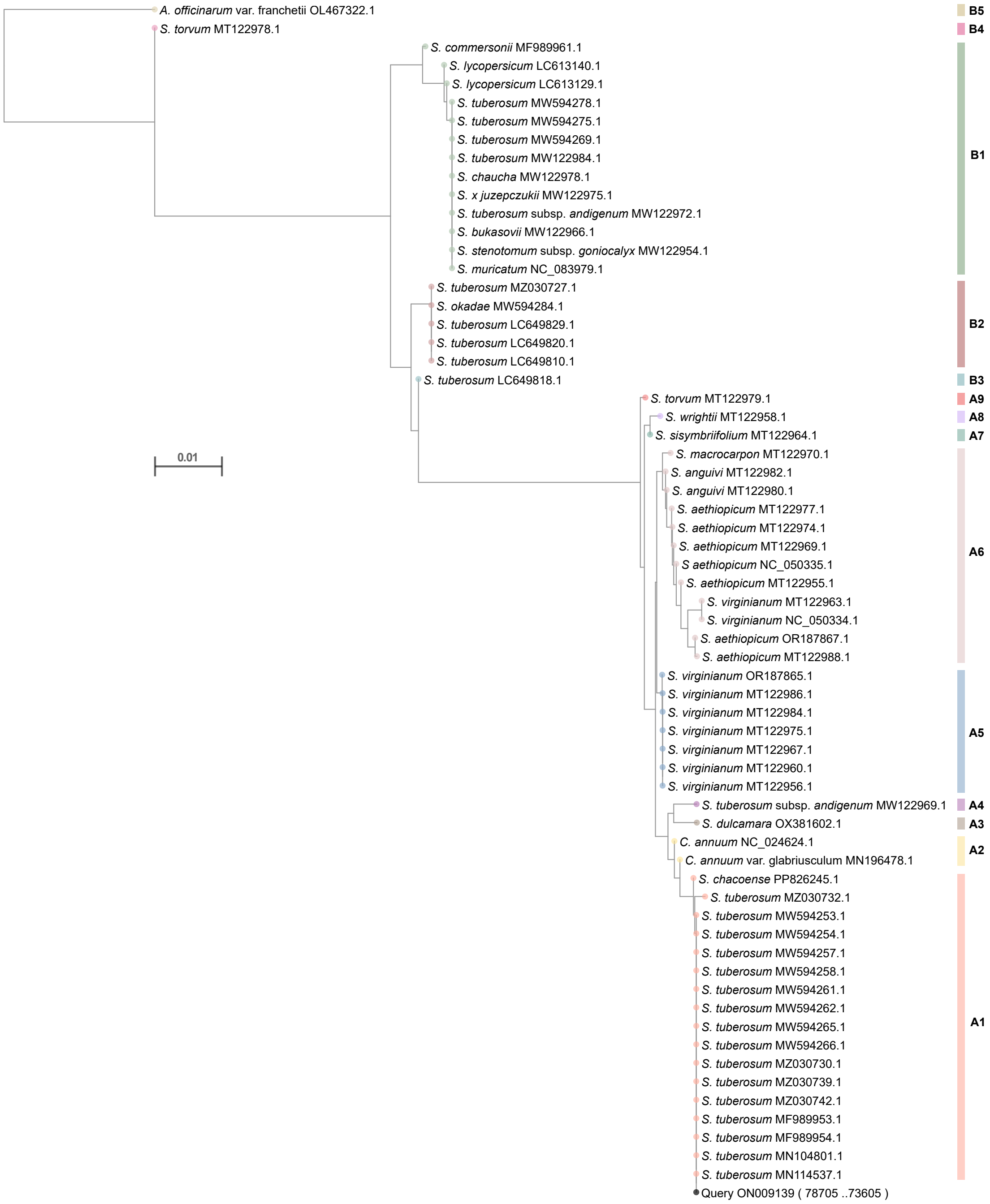


**Figure S6.** Putative groups of accessions obtained by BLAST analysis with the SH9B *orf247-nad4* sequence (P11-P3 primers, see Table S5) as query and Neighbor Joining Method.

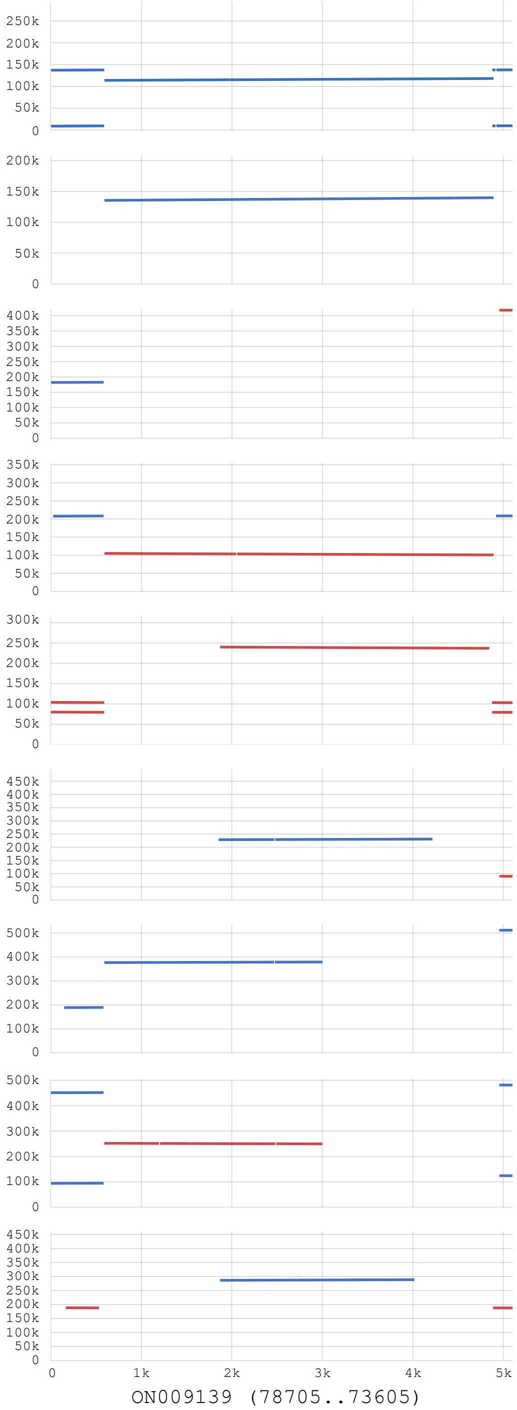


**S. wrightii NN9 (MT122958)**

| Q.start | Q.end | S.start | S.end | %Identity | Length |
|---|---|---|---|---|---|
| 591 | 4892 | 113844 | 118150 | 99.0 | 4307 |
| 1 | 588 | 9041 | 9628 | 95.6 | 588 |
| 1 | 588 | 137159 | 137746 | 95.6 | 588 |
| 4878 | 5101 | 9630 | 9844 | 92.9 | 224 |
| 4878 | 5101 | 137748 | 137962 | 92.9 | 224 |

**S. torvum NN12 (MT122979)**

| Q.start | Q.end | S.start | S.end | %Identity | Length |
|---|---|---|---|---|---|
| 591 | 4892 | 135537 | 139843 | 98.9 | 4307 |

**S. torvum NN12 (MT122978)**

| Q.start | Q.end | S.start | S.end | %Identity | Length |
|---|---|---|---|---|---|
| 1 | 582 | 182061 | 182642 | 94.3 | 582 |
| 4956 | 5101 | 418147 | 418002 | 99.3 | 146 |

**S. sisymbriifolium NN15 (MT122964)**

| Q.start | Q.end | S.start | S.end | %Identity | Length |
|---|---|---|---|---|---|
| 591 | 4892 | 104978 | 100681 | 98.8 | 4307 |
| 23 | 581 | 207776 | 208334 | 94.5 | 559 |
| 4919 | 5101 | 208364 | 208546 | 98.4 | 183 |

**S. tuberosum subsp. Andigenum (MW122969)**

| Q.start | Q.end | S.start | S.end | %Identity | Length |
|---|---|---|---|---|---|
| 1870 | 4845 | 239996 | 237016 | 98.9 | 2981 |
| 1 | 590 | 80072 | 79483 | 100.0 | 590 |
| 1 | 590 | 104027 | 103438 | 100.0 | 590 |
| 4876 | 5101 | 79485 | 79260 | 100.0 | 226 |
| 4876 | 5101 | 103440 | 103215 | 100.0 | 226 |

**C. annuum var. glabriusculum (MN196478)**

| Q.start | Q.end | S.start | S.end | %Identity | Length |
|---|---|---|---|---|---|
| 1854 | 4217 | 228551 | 230905 | 99.5 | 2364 |
| 4958 | 5101 | 90125 | 89982 | 100.0 | 144 |

**S. aethiopicum cv. Aculeatum (OR187867)**

| Q.start | Q.end | S.start | S.end | %Identity | Length |
|---|---|---|---|---|---|
| 591 | 3001 | 376977 | 379383 | 98.7 | 2416 |
| 147 | 582 | 188870 | 189305 | 93.8 | 436 |
| 4956 | 5101 | 512163 | 512308 | 99.3 | 146 |

**S. virginianum cv. EP28 (OR187865)**

| Q.start | Q.end | S.start | S.end | %Identity | Length |
|---|---|---|---|---|---|
| 591 | 3001 | 251926 | 249525 | 98.1 | 2421 |
| 1 | 582 | 93912 | 94493 | 93.3 | 582 |
| 1 | 582 | 451346 | 451927 | 93.3 | 582 |
| 4956 | 5101 | 124022 | 124167 | 99.3 | 146 |
| 4956 | 5101 | 481456 | 481601 | 99.3 | 146 |

**S. dulcamara (OX381602)**

| Q.start | Q.end | S.start | S.end | %Identity | Length |
|---|---|---|---|---|---|
| 1870 | 4017 | 286500 | 288651 | 99.0 | 2152 |
| 167 | 532 | 188530 | 188165 | 93.2 | 366 |
| 4888 | 5101 | 188172 | 187959 | 99.1 | 214 |

**Figure S7.** Dot Plots between the sequence of the "*orf247-nad4*" SH9B region (corresponding to the fragment between nucleotides 78705 and 73605 in ON009139, amplified by P11-P3 primers and including *orf125*, see Table S5) and corresponding sequences identified in some GenBank accessions. The x-and y-axes report the SH9B and corresponding sequences, respectively. Plus/plus matches are in blue, whereas plus/minus matches are in orange.

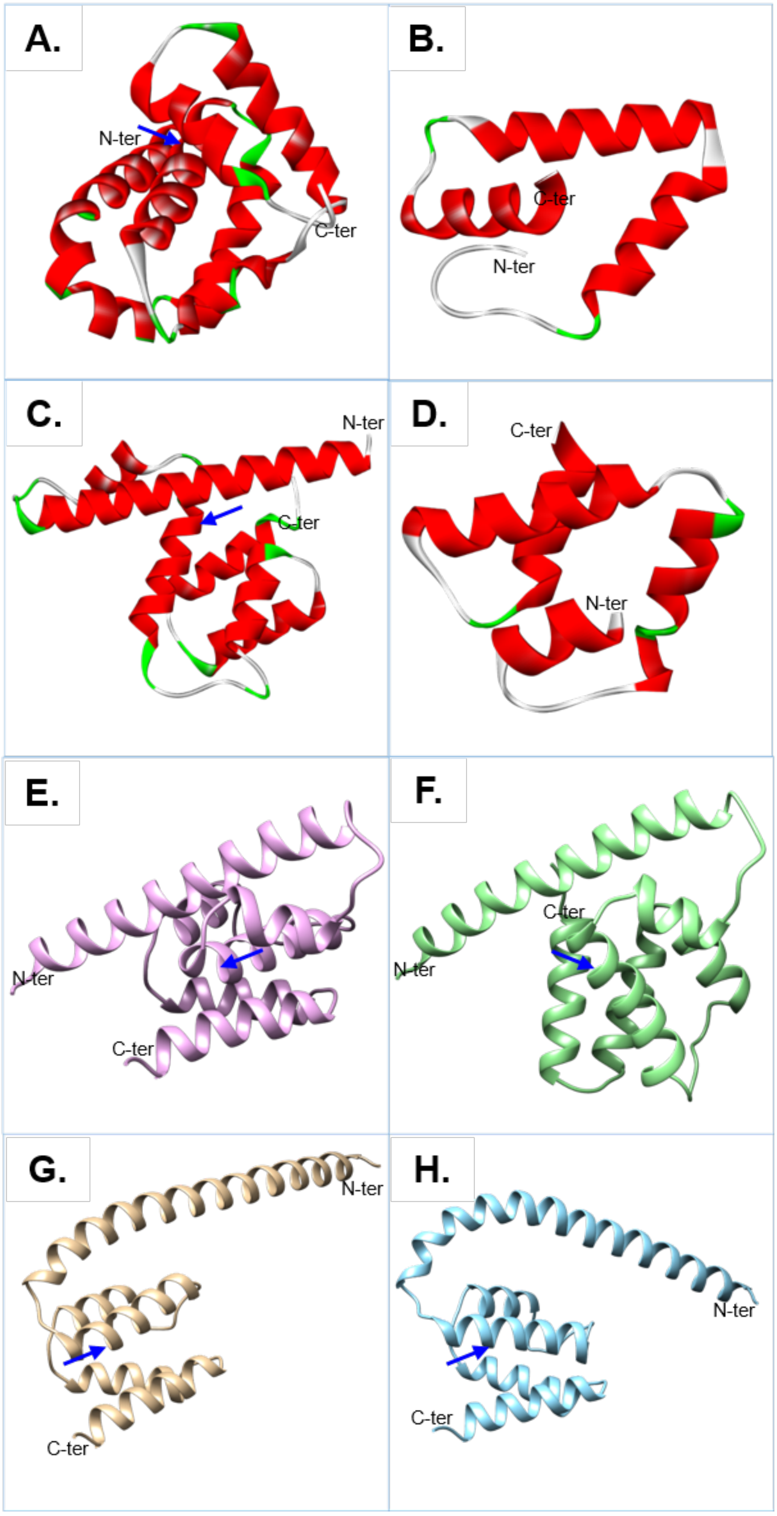


**Figure S8.** Models of ORF125 from *S. tuberosum* Group *Tuberosum* (= SH9B) obtained by various predictors (A–E) and of mutant forms by AlphaFold (F-H). Position 55 is pointed by the blue arrow position, when included in the model. A. Model obtained by I-TASSER of the complete sequence; it is the first in the top ten models produced by the predictor and shows a Z-score equal to -5.22. B. The best model obtained by SWISS MODEL; it covers only a portion of the sequence from Leu64 to Lys119 and shows a Z-score equal to -5.9. C. Complete model of the protein obtained by AlphaFold with a Z-score of -4.72. D. Model obtained by Phyre2 of the ORF125 region from Ile59 to Trp121; it is at the eighth place in the top ten models produced by the predictor but reaches the best Z-score, with a value equal to -6.28. E. Same as C: in this case model is rotated by almost 180 degrees to allow a better comparison to the models presented in the subsequent panels. F. The best model obtained for the edited ORF125 with the D55N mutation, the model shows a Z-score equal to -3.87. G. The complete model of the protein ORF125 from *S. tuberosum* Group *Andigenum* (*adg1)* with a Z-score of -3.46. H. Model of ORF125 from *S. wrightii/sisymbriifolium/torvum;* it reaches a Z-score with a value equal to -3.46.

```
SH9B / tuberosum (wt)    MNIFDIFTIAYSMIFFVSFTFFVARHKLSSYLSFNDLLTREEYLRLLVPEALQKDIQLIL   60
SH9B (D55N)              MNIFDIFTIAYSMIFFVSFTFFVARHKLSSYLSFNDLLTREEYLRLLVPEALQKNIQLIL   60
andigenum                MNILDIFTIAYSMIFFVSFTFFVAVHKLSSYLSFNDLLTREEHLRLLVPEALQKDIQQIL   60
wrightii                 MNILDIFTIAYSMIFFVSFTFFVAGHKLSSYLSFNDLLTREEHLRLLVPEALQKDIQLIL   60
sisymbriifolium          MNILDIFTIAYSMIFFVSFTFFVAGHKLSSYLSFNDLLTREEHLRLLVPEALQKDIQLIL   60
torvum                   MNILDIFTIAYSMIFFVSFTFFVAGHKLSSYLSFNDLLTREEHLRLLVPEALQKDIQLIL   60
                         ***:******************** *****************:***********:** **

SH9B / tuberosum (wt)    NLYLTNLGERMPQGHSWQELAHKIHEGSQDKEFLLSIISDIVSNGMSSEWVLMALQILKD   120
SH9B (D55N)              NLYLTNLGERMPQGHSWQELAHKIHEGSQDKEFLLSIISDIVSNGMSSEWVLMALQILKD   120
andigenum                NLYLNNLGERMPQGHSWQELAHKIHEGSQDKEFLLSIFSDIIYNGTSSEWVLMALQILKD   120
wrightii                 NLYLNNLGERMPQGHSWQELAHKIHEGSQDKEFLLSILSDIIYNGTSSEWVLMALQILKD   120
sisymbriifolium          NLYLNNLGERMPQGHSWQELAHKIHEGSQDKEFLLSILSDIIYNGTSSEWVLMALQILKD   120
torvum                   NLYLNNLGERMPQGHSWQELAHKIHEGSQDKEFLLSILSDIIYNGTSSEWVLMALQILKD   120
                         ****.********************************:***: ** **************

SH9B / tuberosum (wt)    WTFPL*   125
SH9B (D55N)              WTFPL*   125
andigenum                WTFPP*   125
wrightii                 WTFPP*   125
sisymbriifolium          WTFPP*   125
torvum                   WTFPP*   125
                         **** *
```

**Figure S9.** Alignments of ORF125 from SH9B (ON009139) / *cv.* Désirée (MN104801), edited SH9B (Nicolia et al., 2024), *S. tuberosum* Group *Andigenum* (MW122969), *S. wrightii* (MT122958), *S. sisymbriifolium* (MT122964), *S. torvum* (MT122979).

**ORF125$^{1-125}$**

| | D55N | ADG | WRI |
|---|---|---|---|
| **wt** | 10.01 | 8.18 | 9.24 |
| **D55N** | | 17.29 | 15.23 |
| **ADG** | | | 3.77 |

**ORF125$^{1-41}$**

| | D55N | ADG | WRI |
|---|---|---|---|
| **wt** | 1.11 | 0.76 | 0.55 |
| **D55N** | | 1.03 | 0.97 |
| **ADG** | | | 0.26 |

**ORF125$^{42-125}$**

| | D55N | ADG | WRI |
|---|---|---|---|
| **wt** | 1.71 | 3.02 | 3.65 |
| **D55N** | | 2.76 | 3.70 |
| **ADG** | | | 3.12 |

**Figure S10.** Root Mean Square Deviation (RMSD) values derived by the comparison of wt (*tbr*) and mutant forms of the entire ORF125 protein or the N and C-terminal regions (from aminoacids 1 to 41 and from 42 to 125, respectively).

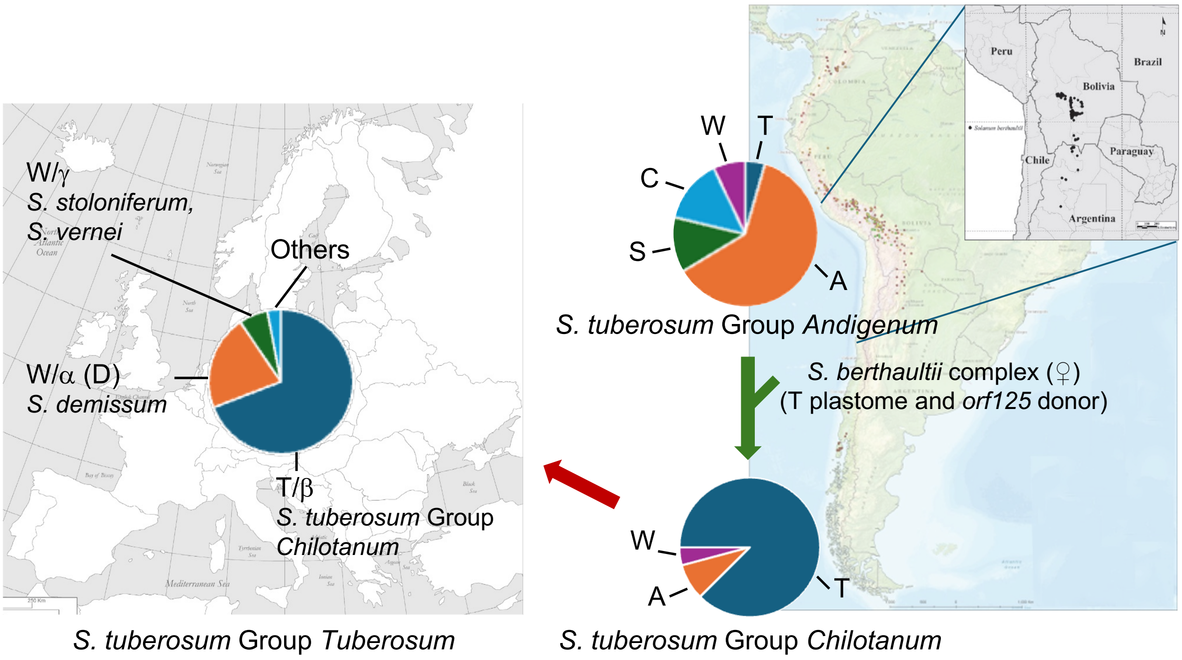


**Figure S11.** Cytoplasm distribution in Andean, Chilean and European cultivated tetraploid potato accessions and putative role of *orf125* in common potato evolution. Within the *berthaultii* complex in South Bolivia and North Argentina (inset), accessions crossed as female with tetraploid *Andigenum* species contributing not only the T-type plastomes but also the chondriomes containing the CMS-inducing *orf125*. This genetic exchange led to the emergence of the new species within the *Chilotanum* group. The novel T/β-cytoplasm subsequently spread to Europe and other parts of the world. Its prevalence in modern cultivars across various collections varies, depending on the use of wild species in breeding programs (Hosaka and Sanetomo, 2012). Data of the *Andigenum* and *Chilotanum* Groups are sourced from Hosaka and Hanneman (Hosaka and Hanneman, 1988) (with European and American cultivars omitted), while information on European cultivars is from Sanetomo and Gebhardt (Sanetomo and Gebhardt, 2015). Modified from Peralta et al. (2021) and Spooner et al. (2007).

**Table S1.** Comparison of the mitochondrial genomes of the somatic hybrids SH9A and SH9B.

| | SH9A | | | | SH9B | | |
|---|---|---|---|---|---|---|---|
| Chromosomes | 1 | 2 | 3 | 4 | 1 | 2 | 3 |
| GenBank acc. no. | ON682437 | ON682438 | ON682439 | ON682440 | ON009139 | ON009140 | ON009141 |
| Genome length (bp) | 251363 | 109928 | 49622 | 48445 | 313767 | 111810 | 48452 |
| Total genes[a] | 94 (109) | | | | 101 (118) | | |
| Protein coding genes[a] | 37 (41) | | | | 37 (41) | | |
| Pseudogenes | 1 | | | | 2 | | |
| Hypothetical genes[a] | 35 (41) | | | | 41 (49) | | |
| rRNA[a] | 3 (5) | | | | 3 (5) | | |
| tRNA[a] | 18 (21) | | | | 18 (21) | | |
| Total genes[ab] | 55 (62) | 22 (23) | 5 (11) | 12 (13) | 65 (80) | 24 (25) | 12 (13) |
| Protein coding genes[ab] | 19 | 12 | 1 (5) | 5 | 20 (24) | 12 | 5 |
| Pseudogenes[b] | 0 | 1 | 0 | 0 | 0 | 2 | 0 |
| Hypothetical genes[ab] | 25 (30) | 6 | 1 (2) | 3 | 31 (39) | 7 | 3 |
| rRNA[ab] | 3 (5) | 0 | 0 | 0 | 3 (5) | 0 | 0 |
| tRNA[ab] | 8 | 3 (4) | 3 (4) | 4 (5) | 11 (12) | 3 (4) | 4 (5) |
| GC%[b] | 45.40 | 44.56 | 44.58 | 44.59 | 45.19 | 44.59 | 44.59 |
| AT/GC[b] | 1.20 | 1.24 | 1.24 | 1.24 | 1.21 | 1.24 | 1.24 |

[a] First value excludes duplicated; value in parentheses includes them.

[b] Values referring to individual chromosomes

**Table S2.** Details on the syntenic building blocks shared between somatic hybrids and their parental species. The following information is associated with each syntenic block: block identifier, chromosome accession number, strand, start and end positions, length of the block, list of genes included in the block, and any notes.

| SyntenyBlocks | SeqID[a] | Strand | Start | End | Length | Genes | Notes |
|---|---|---|---|---|---|---|---|
| 14 | MF989961 | + | 1 | 85223 | 85223 | *trnP-UGG, trnF-GAA, trnS-GCU, orfx, orf140, rrn26, trnfM-CAU, nad2, trnY-GUA, trnN-GUU, trnC-GCA, nad5, nad4, orf247*, rps4, nad6, nad4L, atp4* | orfx is annotated as mttb in (MN104801, ON682437, ON009139), orf140 overlaps orf265a annotated in (MN104801, ON682437, ON009139), orf125a is absent in assembled genome, orf161a is absent in assembled genome |
| | MN104801 | + | 15473 | 104990 | 89518 | *trnP-UGG, trnF-GAA, trnS-GCU, mttb, orf265a, rrn26, trnfM-CAU, nad2, trnY-GUA, trnN-GUU, trnC-GCA, nad5, nad4, orf125a, orf161a, orf247, rps4, nad6, nad4L, atp4* | mttb is annotated as orfx in MF989961, orf265a overlaps orf140 annotated in MF989961 |
| | ON682437 | + | 131192 | 216863 | 85672 | *trnP-UGG, trnF-GAA, trnS-GCU, mttb, orf265a, rrn26, trnfM-CAU, nad2, trnY-GUA, trnN-GUU, trnC-GCA, nad5, nad4, orf247, rps4, nad6, nad4L, atp4* | mttb is annotated as orfx in MF989961, orf125a is absent in assembled genome, orf161a is absent in assembled genome |
| | ON009139 | + | 15492 | 105461 | 89970 | *trnP-UGG, trnF-GAA, trnS-GCU, mttb, orf265a, rrn26, trnfM-CAU, nad2, trnY-GUA, trnN-GUU, trnC-GCA, nad5, nad4, orf125a, orf161a, orf247, rps4, nad6, nad4L, atp4* | mttb is annotated as orfx in MF989961, orf265a overlaps orf140 annotated in MF989961 |
| 10 | MF989960 | + | 72589 | 128725 | 56137 | *ccmFc, orf169, orf102, trnN-GUU, cob, trnS-UGA, orf123, trnD-GUC, trnS-GGA, sdh4, cox3, orfB, orf118*, orf152* | orf102 is annotated as orf106 in (MN104802, ON682438, ON009140), orfB is annotated as atp8 in (MN104802, ON682438, ON009140), orf152 partially overlaps rps1 annotated in (MN104802, ON682438, ON009140) |
| | MN104802 | + | 2854 | 59112 | 56259 | *ccmFC, orf169, orf106, trnN-GUU, cob, trnS-UGA, orf123, trnD-GUC, trnS-GGA, sdh4, cox3, atp8, orf118, rps1* | orf106 is annotated as orf102 in MF989960, atp8 is annotated as orfB in MF989960, orf52 is not annotated |
| | ON682438 | + | 2854 | 58992 | 56139 | *ccmFC, orf169, orf106, trnN-GUU, cob, trnS-UGA, orf123, trnD-GUC, trnS-GGA, sdh4, cox3, atp8, orf118, rps1* | orf106 is annotated as orf102 in MF989960, atp8 is annotated as orfB in MF989960, orf52 is not annotated |
| | ON009140 | + | 2854 | 59062 | 56209 | *ccmFC, orf169, orf106, trnN-GUU, cob, trnS-UGA, orf123, trnD-GUC, trnS-GGA, sdh4, cox3, atp8, orf118, rps1* | orf106 is annotated as orf102 in MF989960, atp8 is annotated as orfB in MF989960, orf52 is not annotated |
| 5 | MF989960 | + | 200505 | 239145 | 38641 | *rpl10, rpl2, trnE-UUC, orf103, sdh3, nad2, trnW-CCA, trnP-UGG, nad9, trnH-GUG* | orf210 is absent (partial sequence found) in assembled genome, orf103 is annotated as orf96 in (MN104803,ON682440, ON009141) |
| | MF989961 | + | 147829 | 186469 | 38641 | *rpl10, rpl2, trnE-UUC, orf103, sdh3, nad2, trnW-CCA, trnP-UGG, nad9, trnH-GUG* | orf210 is absent (partial sequence found) in assembled genome, orf103 is annotated as orf96 in (MN104803,ON682440, ON009141) |
| | MN104803 | + | 10588 | 49230 | 38643 | *rpl10, orf210, rpl2, trnE-UUC, orf96, sdh3, nad2, trnW-CCA, trnP-UGG, nad9, trnH-GUG* | orf96 is annotated as orf 103 in (MF989960, MF989961) |
| | ON682440 | + | 10588 | 48445 | 37858 | *rpl10, rpl2, trnE-UUC, orf96, sdh3, nad2, trnW-CCA, trnP-UGG, nad9, trnH-GUG* | orf210 is absent (partial sequence found) in assembled genome, orf96 is annotated as orf103 in (MF989960, MF989961) |
| | ON009141 | + | 10593 | 48452 | 37860 | *rpl10, rpl2, trnE-UUC, orf96, sdh3, nad2, trnW-CCA, trnP-UGG, nad9, trnH-GUG* | orf210 is absent in assembled genome, orf96 is annotated as orf103 in (MF989960, MF989961) |
| 21 | MN104802 | + | 83719 | 111586 | 27868 | *rps12, nad3, orf265b* | |
| | ON682438 | + | 81763 | 109647 | 27885 | *rps12, nad3, orf265b* | |
| | ON009140 | + | 83684 | 111529 | 27846 | *rps12, nad3, orf265b* | |
| 9 | MF989960 | + | 161701 | 188790 | 27090 | *orf103*, nad7, orf109, orf125, trnM-CAU* | |
| | MF989961 | + | 109025 | 136114 | 27090 | *orf103*, nad7, orf109, orf125, trnM-CAU* | |
| | MN104801 | - | 285257 | 258173 | 27085 | *orf103, nad7, orf109, orf125c, trnI-CAU* | |
| | ON682437 | - | 112303 | 85219 | 27085 | *orf103, nad7, orf109, orf125c, trnI-CAU* | |
| | ON009139 | - | 288429 | 261345 | 27085 | *orf103, nad7, orf109, orf125c, trnI-CAU* | |
| 17 | MF989960 | + | 1 | 22932 | 22932 | *trnM-CAU, orf104, trnG-GCC, trnQ-UUG* | orf104 is annotated as orf126 in (MN104801, ON682439, ON009139) |
| | MN104801 | + | 184552 | 207484 | 22933 | *trnM-CAU, orf126, trnG-GCC, trnQ-UUG* | orf126 is annotated as orf104 in MF989960 |
| | ON682439 | + | 26685 | 49622 | 22938 | *trnM-CAU, orf126, trnG-GCC, trnQ-UUG* | orf126 is annotated as orf104 in MF989960 |
| | ON009139 | + | 187715 | 210646 | 22932 | *trnM-CAU, orf126, trnG-GCC, trnQ-UUG* | orf126 is annotated as orf104 in MF989960 |
| 13 | MF989960 | + | 30435 | 49807 | 19373 | *orf125, orf77*, nad1* | |
| | MN104801 | + | 214987 | 234293 | 19307 | *orf125b, orf77, nad1* | |
| | ON682437 | - | 51267 | 31961 | 19307 | *orf125b, orf77, nad1* | |
| | ON009139 | + | 218149 | 237455 | 19307 | *orf125b, orf77, nad1* | |
| 12 | MF989960 | + | 280637 | 298596 | 17960 | *orf159*, orf152*, nad5, orf122, matR* | orf105b is not annotated: presents 1 gap (premature stop codon) compared to orf105b annotated in (MN104801, ON009139) |
| | MN104801 | - | 137041 | 119114 | 17928 | *orf159, orf105b, orf152, nad5, orf122a, matR* | |
| | ON682437 | - | 250380 | 232436 | 17945 | *orf159, orf152, nad5, orf122a, matR* | orf105b is not annotated: presents 1 gap (premature stop codon) compared to orf105b annotated in (MN104801, ON009139) |

### Table S2. (cont.d)

| | SyntenyBlocks | SeqID[a] | Strand | Start | End | Length | Genes | Notes |
|---|---|---|---|---|---|---|---|---|
| | | ON009139 | - | 138896 | 121038 | 17859 | *orf159, orf105b, orf152, nad5, orf122a, matR* | |
| | 15 | MF989960 | + | 322688 | 338427 | 15740 | *trnC-GCA, ccmC* | |
| | | MN104801 | + | 169872 | 184551 | 14680 | *trnC-GCA, ccmC* | |
| | | ON682439 | + | 11460 | 26684 | 15225 | *trnC-GCA, ccmC* | |
| | | ON009139 | + | 171689 | 187714 | 16026 | *trnC-GCA, ccmC* | |
| | 1 | MF989960 | - | 152023 | 136449 | 15575 | *orf304*, orf102, rrn18, orf108, rrn5, orf100, orf240, orf101* | orf100 overlaps orf141 annotated in (MN104801,ON682437,ON009139), orf240 overlaps orf438 annotated in (MN104801,ON682437,ON009139) |
| | | MF989960 | + | 298597 | 314171 | 15575 | *orf304*, orf102, rrn18, orf108, rrn5, orf100, orf438*, orf101* | orf100 overlaps orf141 annotated in (MN104801,ON682437,ON009139) |
| | | MF989961 | - | 99347 | 85224 | 14124 | *orf304*, orf102, rrn18, orf108, rrn5, orf100, orf240, orf101* | orf100 overlaps orf141 annotated in (MN104801,ON682437,ON009139), orf240 overlaps orf438 annotated in (MN104801,ON682437,ON009139) |
| | | MN104801 | - | 119113 | 104991 | 14123 | *orf304, orf102a, rrn18, orf108, rrn5, orf141, orf438, orf101* | orf141 overlaps orf100 annotated in (MF989960, MF989961), orf438 overlaps orf240 annotated in (MF989960, MF989961) |
| | | MN104801 | + | 294931 | 310479 | 15549 | *orf304 Ψ, orf102a, rrn18, orf108, rrn5, orf141, orf438, orf101* | orf141 overlaps orf100 annotated in (MF989960, MF989961), orf438 overlaps orf240 annotated in (MF989960, MF989961) |
| | | ON682437 | - | 15619 | 36 | 15584 | *orf304, orf102a, rrn18, orf108, rrn5, orf141, orf438, orf101* | orf141 overlaps orf100 annotated in (MF989960, MF989961), orf438 overlaps orf240 annotated in (MF989960, MF989961) |
| | | ON682437 | - | 232435 | 216864 | 15572 | *orf102a, rrn18, orf108, rrn5, orf141, orf438, orf101* | orf304 is abstent (3 gaps - premature stop codon), orf141 overlaps orf100 annotated in (MF989960, MF989961), orf438 overlaps orf240 annotated in (MF989960, MF989961) |
| | | ON009139 | + | 298110 | 313685 | 15576 | *orf304, orf102a, rrn18, orf108, rrn5, orf141, orf438, orf101* | orf141 overlaps orf100 annotated in (MF989960, MF989961), orf438 overlaps orf240 annotated in (MF989960, MF989961) |
| | | ON009139 | - | 121037 | 105462 | 15576 | *orf304, orf102a, rrn18, orf108, rrn5, orf141, orf438, orf101* | orf141 overlaps orf100 annotated in (MF989960, MF989961), orf438 overlaps orf240 annotated in (MF989960, MF989961) |
| | 16 | MF989960 | + | 266744 | 279653 | 12910 | *ccmFN, orf105, cox1, rps10, rps14 Ψ*, rpl5* | orf122c is absent in assembled genome, cob Ψ is abstent in assembled genome |
| | | MN104802 | + | 68973 | 83718 | 14746 | *ccmFN, orf105c, cox1, rps10, orf122c, cob Ψ, rps14 Ψ, rpl5* | |
| | | ON682438 | + | 68853 | 81762 | 12910 | *ccmFN, orf105c, cox1, rps10, rps14 Ψ, rpl5* | orf122c is absent in assembled genome, cob Ψ is abstent in assembled genome |
| | | ON009140 | + | 68923 | 83683 | 14761 | *ccmFN, orf105c, cox1, rps10, orf122c, cob Ψ, rps14 Ψ, rpl5* | |
| | 7 | MF989960 | + | 239146 | 249740 | 10595 | *orf109, trnK-UUU, orf103, ccmB* | orf109 is annotated as orf109d in MF989961 and orf116 in (MN104803, ON682440, ON009141), orf103 is annotated as orf113 in (MN104803, ON682440, ON009141) |
| | | MF989961 | + | 186470 | 197064 | 10595 | *orf109d, trnK-UUU, orf113*, ccmB* | orf109d is annotated as orf109 in MF989960 and orf116 in (MN104803, ON682440, ON009141) |
| | | MN104803 | + | 1 | 10587 | 10587 | *orf116, trnK-UUU, orf113, ccmB* | orf116 is annotated as orf109 in MF989960 and orf109d in MF989961, orf113 is annotated as orf103 in MF989960 |
| | | ON682440 | + | 1 | 10587 | 10587 | *orf116, trnK-UUU, orf113, ccmB* | orf116 is annotated as orf109 in MF989960 and orf109d in MF989961, orf113 is annotated as orf103 in MF989960 |
| | | ON009141 | + | 1 | 10592 | 10592 | *orf116, trnK-UUU, orf113, ccmB* | orf116 is annotated as orf109 in MF989960 and orf109d in MF989961, orf113 is annotated as orf103 in MF989960 |
| | 6 | MF989960 | + | 152024 | 161700 | 9677 | *rps13, nad1, orf110* | orf110 is annotated as orf122b in (MN104801,ON682437,ON009139) |
| | | MF989961 | + | 99348 | 109024 | 9677 | *rps13, nad1, orf110* | orf110 is annotated as orf122b in (MN104801,ON682437,ON009139) |
| | | MN104801 | - | 294930 | 285258 | 9673 | *rps13, nad1, orf122b* | orf122b is annotated as orf110 in (MF989960, MF989961) |
| | | ON682437 | + | 15620 | 25243 | 9624 | *rps13, nad1, orf122b* | orf122b is annotated as orf110 in (MF989960, MF989961) |
| | | ON009139 | - | 298109 | 288430 | 9680 | *rps13, nad1, orf122b* | orf122b is annotated as orf110 in (MF989960, MF989961) |
| | 20 | MN104802 | + | 59113 | 68972 | 9860 | | |
| | | ON682438 | + | 58993 | 68852 | 9860 | | |
| | | ON009140 | + | 59063 | 68922 | 9860 | | |
| | 8 | MF989960 | + | 257683 | 266352 | 8670 | *orf111, orf185, orf119*, atp1* | |
| | | MF989961 | + | 205007 | 213676 | 8670 | *orf111, orf185, orf119*, atp1* | |
| | | MN104801 | + | 6807 | 15472 | 8666 | *orf185, orf119, atp1* | orf111 is absent in assembled genome ( 1 gap - premature stop codon) |
| | | ON682437 | + | 122515 | 131191 | 8677 | *orf111, orf185, orf119, atp1* | |

### Table S2. (cont.d)

| | SyntenyBlocks | SeqID[a] | Strand | Start | End | Length | Genes | Notes |
|---|---|---|---|---|---|---|---|---|
| | | ON009139 | + | 6815 | 15491 | 8677 | *orf111, orf185, orf119, atp1* | |
| | 18 | MN104801 | - | 145700 | 137042 | 8659 | *orf137* | |
| | | ON682437 | + | 71397 | 80077 | 8681 | *orf137* | |
| | | ON009139 | - | 147578 | 138920 | 8659 | *orf137* | |
| | 23 | MN104801 | + | 250600 | 258172 | 7573 | | |
| | | ON009139 | + | 253772 | 261344 | 7573 | | |
| | 2 | MF989960 | + | 22933 | 30434 | 7502 | *rps19, rps3, orf102, rpl16, cox2* | |
| | | MN104801 | + | 158328 | 165829 | 7502 | *rps19, rps3, orf102b, rpl16, cox2* | |
| | | MN104801 | + | 207485 | 214986 | 7502 | *rps19, rps3, orf102b, rpl16, cox2* | |
| | | ON682437 | - | 58769 | 51268 | 7502 | *rps19, rps3, orf102b, rpl16, cox2* | |
| | | ON682439 | + | 1 | 7502 | 7502 | *rps19, rps3, orf102b, rpl16, cox2* | |
| | | ON009139 | + | 160206 | 167707 | 7502 | *rps19, rps3, orf102b, rpl16, cox2* | |
| | | ON009139 | + | 210647 | 218148 | 7502 | *rps19, rps3, orf102b, rpl16, cox2* | |
| | 4 | MF989960 | + | 249741 | 257128 | 7388 | *nad5, orf185, orf138* | orf185 is annotated as orf161b in (MN104801, ON682437, ON009139), orf138 in annotated as orf147 in (MN104801, ON682437, ON009139) |
| | | MF989961 | + | 197065 | 204452 | 7388 | *nad5, orf185, orf138* | orf185 is annotated as orf161b in (MN104801, ON682437, ON009139), orf138 in annotated as orf147 in (MN104801, ON682437, ON009139) |
| | | MN104801 | + | 243216 | 250599 | 7384 | *nad5, orf161b, orf147* | orf161b is annotated as orf185 in (MF989960, MF989961), orf147 is annotated as orf138 in (MF989960, MF989961) |
| | | ON682437 | + | 114573 | 121960 | 7388 | *nad5, orf161b, orf147* | orf161b is annotated as orf185 in (MF989960, MF989961), orf147 is annotated as orf138 in (MF989960, MF989961) |
| | | ON009139 | + | 246388 | 253771 | 7384 | *nad5, orf161b, orf147* | orf161b is annotated as orf185 in (MF989960, MF989961), orf147 is annotated as orf138 in (MF989960, MF989961) |
| | 22 | MN104801 | + | 1 | 6806 | 6806 | *orf77, nad1, orf146, orf105a* | |
| | | ON009139 | + | 1 | 6814 | 6814 | *orf77, nad1, orf146, orf105a* | |
| | 3 | MF989960 | + | 191579 | 198289 | 6711 | *orf263 cox2-like** | |
| | | MF989961 | + | 138903 | 145613 | 6711 | *orf263 cox2-like** | |
| | | MN104801 | + | 234294 | 241000 | 6707 | *orf263 cox2-like* | |
| | | ON682437 | - | 31960 | 25244 | 6717 | *orf263 cox2-like* | |
| | | ON009139 | + | 237456 | 244172 | 6717 | *orf263 cox2-like* | |
| | 11 | MF989960 | + | 66177 | 72588 | 6412 | *nad1* | |
| | | MN104801 | + | 145701 | 152319 | 6619 | *nad1* | |
| | | ON682437 | - | 71396 | 64778 | 6619 | *nad1* | |
| | | ON009139 | + | 147579 | 154197 | 6619 | *nad1* | |
| | 19 | MN104801 | - | 158327 | 152320 | 6008 | *orf509* | |
| | | ON682437 | + | 58770 | 64777 | 6008 | | orf509 is absent (truncated at 5') |
| | | ON009139 | - | 160205 | 154198 | 6008 | | orf509 is absent (truncated at 5') |

[a] MF989960, MF989961 (*Solanum commersonii*); MN104801, MN104802, MN104803 (*S. tuberosum*); ON682437, ON682438, ON682439, ON682440 (SH9A); ON009139, ON009140, ON009141 (SH9B)

* Present but not annotated

**Table S3.** Pollen production and stainability in regenerated (V0) and tuber-derived propagated plants (V1). The type of mutations induced by editing and their homoplasmy/heteroplasmy status is also indicated.

| Clone | V0 plants | | | V1 plants | | |
|---|---|---|---|---|---|---|
| | Genotype[a] | Pollen production[b] | Pollen stainability (%) | Genotype | Pollen production | Pollen stainability (%) |
| *A) mitoTALEN* | | | | | | |
| T1-6 | Del1066 (het.) | + / +++ | 100 | Del1066 (het.) | ++ / +++ | 100 |
| T1-29 | Del1066 (het.) | ++ / +++ | 100 | Del1066 (het.) | ++ / +++ | 100 |
| T1-49 | Del1066 (het.) | ++ / +++ | 100 | Del1066 (het.) + N[c] | ++ / +++ | 100 |
| T2-1[d] | Del236 (het.) | - | - | Del236 (het.) | na[e] | na |
| T2-3 | WT | - | - | WT | - | - |
| T2-10 | Del236 (hom.) | +++ | 100 | Del236 (hom.) | +++ | 100 |
| T2-11 | WT | - | - | na | na | na |
| T2-12 | Del4288 (hom.) | + / ++ | 100 | Del4288 (hom.) | + / ++ | 100 |
| T2-14 | Del1066 (hom.) | +++ | 100 | Del1066 (hom.) | +++ | 100 |
| T2-15 | WT | - | - | WT | - | - |
| T2-23 | Del236 (het.) | - | - | WT | - | - |
| T2-24 | WT | - | - | na | na | na |
| T2-26 | Del236 (het.) + Ins4[f] | +++ | 100 | WT (Ins4) | +++ | 100 |
| T2-28 | WT | - | - | WT | - | - |
| T2-29[d] | Del236 (het.) | - | - | Del236 (het.) + N | na | na |
| T2-30 | WT | - | - | WT | - | - |
| T2-31 | Del236 (hom.) | +++ | 100 | Del236 (hom.) | +++ | na |
| *B) mitoTALENCD* | | | | | | |
| D1-5 | G163A; C169C/T[g] | +++ | 100 | na | na | na |
| D1-28 | G163A; C169C/T | +++ | 100 | na | na | na |
| D1-48 | G163A; C169C/T | +++ | 70-100 | G163A; C169T | +++ | 100 |
| D1-51 | G163A | +++ | 100 | G163A | +++ | 100 |
| D1-54 | G163A; C169C/T | ++ / +++ | 80-100 | na | na | na |
| D1-57 | G163A; C169C/T | +++ | 100 | na | na | na |
| D1-61 | G163A; C169C/T | +++ | 100 | na | na | na |
| D1-68 | G163A; C169C/T | +++ | 100 | na | +++ | 100 |
| D1-69 | G163A; C169C/T | +++ | 100 | G163A; C169C/T | +++ | 100 |
| D1-70[d] | G163A | +++ | 100 | G163A; C169C/T | +++ | na |
| D1-84 | G163A; C169T | +++ | 100 | G163A; C169T | +++ | 100 |
| D1-85 | G163A | +++ | 100 | na | +++ | na |
| D1-90 | G163A; C169C/T | +++ | 100 | G163A | na | na |
| D1-93 | G163A; C169C/T | +++ | 100 | G163A; C169C/T | +++ | 100 |
| D1-103 | G163A; C169C/T | +++ | 80-100 | na | na | na |
| D1-113[d] | G163G/A | - | - | G163G/A; C169C/T | na | na |
| D2-1 | WT | - | - | na | na | na |
| D2-3 | WT | - | - | na | na | na |
| D2-9 | WT | - | - | WT | - | - |
| D2-13 | WT | - | - | na | na | na |
| D2-16 | WT | - | - | na | na | na |

**Table S3. (cont.d)**

| Clone | V0 plants | | | V1 plants | | |
|---|---|---|---|---|---|---|
| | Genotype[a] | Pollen production[b] | Pollen stainability | Genotype | Pollen production | Pollen stainability |
| D2-17 | WT | - | - | na | na | na |
| D2-18 | WT | - | - | na | na | na |
| D2-19 | G256G/A; G258G/A; G259G/A; C263C/T; C265T | +++ | 100 | na | na | na |
| D2-20 | G256G/A; C263C/T; C265T | +++ | 100 | na | na | na |
| D2-21 | WT | - | - | na | na | na |
| D2-22 | G256G/A; G258G/A; G259G/A; C263C/T; C265T | +++ | 100 | na | na | na |
| D2-23 | C265T | +++ | 100 | C265T | +++ | 100 |
| D2-24 | G256G/A; G258G/A; G259G/A; C263C/T; C265T | ++ | 100 | na | na | na |
| D2-27 | C265C/T | - | - | C265C/T | - | - |
| D2-29 | C263C/T; C265T | +++ | 100 | na | na | na |
| D2-32 | C265C/T | - | - | WT | - | - |
| D2-33 | WT | - | - | WT | - | - |
| D2-34 | C265T | +++ | 100 | C265T | ++ | 100 |
| D2-38 | WT | - | - | WT | - | - |
| *C) Control clones* | | | | | | |
| SH9A | | +++ | 100 | | +++ | 100 |
| SH9B | | - | - | | - | - |

[a] For details see Nicolia et al. (2024)

[b] Multiple plants per clone and multiple flowers per plant were generally analyzed for male fertility in various environments. Pollen production was rated from – (nil) to +++ (very abundant). Different ratings separated by a slash indicate variability between flowers. Pollen grains were stained with acetocarmine

[c] T1-49 and T2-29 clones showed novel bands (N) after propagation

[d] The four V1 clones indicated derived by micropropagation *in vitro*

[e] na, not available

[f] The T2-26 clone, besides being heteroplasmic for the 236 bp deletion, had also an insertion of 4 bp in the "wild-type" amplicon

[g] Heteroplasmic base substitution

**Table S4.** List of *Solanum* spp. genotypes used in this study.

| Species | Accession[a] | Code | Origin | Ploidy | Notes |
|---|---|---|---|---|---|
| *S. berthaultii* | PI 498075 | *ber1* | Bolivia | 2*x* | |
| | PI 498095 | *ber2* | Bolivia | 2*x* | |
| | PI 498101 | *ber3* | Bolivia | 2*x* | |
| *S. brachistotrichum* | PI 320265 | *bst* | Mexico | 2*x* | |
| *S. bulbocastanum* | PI 275187 | *blb* | Mexico | 2*x* | |
| *S. cardiophyllum* | PI 347759 | *cph* | Mexico | 2*x* | |
| *S. chacoense* | PI 320282 | *chc* | Argentina | 2*x* | |
| *S. commersonii* | PI 243503 | *cmm* | Argentina | 2*x* | |
| *S. etuberosum* | PI 558054 | *etb* | Chile | 2*x* | |
| *S. infundibuliforme* | PI 472857 | *ifd* | Argentina | 2*x* | |
| *S. nigrum* | - | *ngr* | - | 6*x* | |
| *S. pinnatisectum* | PI 275236 | *pnt* | Mexico | 2*x* | |
| *S. polytrichon* | PI 184773 | *plt* | Mexico | 4*x* | |
| *S. raphanifolium* | PI 265878 | *rap* | Peru | 2*x* | |
| *S. sanctae-rosae* | PI 218221 | *sct* | Argentina | 2*x* | |
| *S. sparsipilum* | PI 234014 | *spl2* | Bolivia | 2*x* | |
| *S. spegazzinii* | PI 205394 | *spg* | Argentina | 2*x* | |
| *S. tarijense* | PI 265577 | *tar1* | Bolivia | 2*x* | |
| | PI 414148 | *tar2* | Argentina | 2*x* | |
| | PI 442689 | *tar3* | Argentina | 2*x* | |
| *S. trifidum* | PI 255541 | *trf* | Mexico | 2*x* | |
| *S. tuberosum* Group *Andigenum* | CPC 843 | *adg* | - | 4*x* | |
| *S. tuberosum* Group *Tuberosum* | - | Des | - | 4*x* | *cv.* Désirée (Urgenta x Depesche)[b] |
| | DH81-7-1463 | SVP11 | - | 2*x* | Dihaploid clone of W72-22-492 |
| *cmm* (+) SVP11[c] | - | SH1A | - | 4*x* | Somatic hybrid, male sterile |
| *cmm* (+) SVP11 | - | SH7A | - | 4*x* | Somatic hybrid, male sterile |
| *cmm* (+) SVP11 | - | SH9A | - | 4*x* | Somatic hybrid, male fertile |
| *cmm* (+) SVP11 | - | SH9B | - | 4*x* | Somatic hybrid, male sterile |
| *cmm* (+) SVP11 | - | SH12A | - | 4*x* | Somatic hybrid, male sterile |
| *cmm* (+) SVP11 | - | SH25A | - | 6*x* | Somatic hybrid, male sterile |

[a] PI and CPC accessions were kindly provided by Potato Introduction Station, Sturgeon Bay, Wisconsin, and Commonwealth Potato Collection, Dundee, UK, respectively; - = not available/relevant

[b] https://www.plantbreeding.wur.nl/PotatoPedigree/index.html

[c] Cardi et al. (1993)

**Table S5.** List of primers used.

| Primers | Sequence (5'-3') | Tm (°C) | Use |
|---|---|---|---|
| orf125 NcoI F | CATGCCATGGGCATGAATATCTTTGATATTTTC | 76 | *orf125* cloning in plant vectors |
| orf125 BglII R | CTAGATCTGCTAGAGGAAAGGTCCAATCTT | 77 | *orf125* cloning in plant vectors / PCR analysis of transgenic plants |
| P*rbcS* F | CCGTTAGATAGCAAACAACA | 64 | PCR analysis of transgenic plants |
| P*lat52* F | AGGCGCGCCCCTATACCCCTTGGATAA | 81 | *orf125* cloning in plant vectors / PCR analysis of transgenic plants |
| P*lat52* R | CTCTAGATTTAAATTGGAATTTTTTTTTTGG | 69 | *orf125* cloning in plant vectors / PCR analysis of transgenic plants |
| P*ta29* F | AGGCGCGCCAACTGGTCTCAACCTCGTA | 81 | *orf125* cloning in plant vectors / PCR analysis of transgenic plants |
| P*ta29* R | CTCTAGATTTTAGCTAAGTTTATTTAAG | 69 | *orf125* cloning in plant vectors / PCR analysis of transgenic plants |
| RT orf125 F | CGTAGCCAGACACAAACTTTC | 69 | RT-PCR analysis of *orf125* expression in somatic hybrids |
| RT orf125 R | TGCAAAGCCATCAAGACCCA | 68 | RT-PCR analysis of *orf125* expression in somatic hybrids |
| RT 18S F | TAGATAAAAGGTCGACGCGG | 68 | RT-PCR analysis of *rrn18* expression in somatic hybrids |
| RT 18S R | CCCAAAGTCCAACTACGAGC | 70 | RT-PCR analysis of *rrn18* expression in somatic hybrids |
| qRT-PCR orf125 F | CCACAAAATTCACGAGGGCT | 68 | qRT-PCR analysis of *orf125* expression in transgenic plants |
| qRT-PCR orf125 R | AGCCATCAAGACCCATTCACT | 69 | qRT-PCR analysis of *orf125* expression in transgenic plants |
| qRT-PCR ef1α F | ATTGGAAACGGATATGCTCCA | 63 | qRT-PCR analysis of *ef1α* expression in transgenic plants |
| qRT-PCR ef1α R | TCCTTACCTGAACGCCTGTCA | 66 | qRT-PCR analysis of *ef1α* expression in transgenic plants |
| P4 | TTATAGAGGAAAGGTCCAATCTTICA | 62 | PCR amplification of *orf125* coding sequence |
| P5 | ATGAATATCTTTGATATTTTCACG | 63 | PCR amplification of *orf125* coding sequence |
| P3 | GCACGGGACAAAGAATAAGACC | 62 | PCR amplification of *orf247-nad4* genomic region |
| P11 | TTATTATTCGGGCGGGGCTC | 60 | PCR amplification of *orf247-nad4* genomic region |
